# SUSD4 controls GLUA2 degradation, synaptic plasticity and motor learning

**DOI:** 10.1101/859587

**Authors:** I. González-Calvo, K. Iyer, M. Carquin, A. Khayachi, F.A. Giuliani, J. Vincent, M. Séveno, S.M. Sigoillot, M. Veleanu, S. Tahraoui, M. Albert, O. Vigy, C. Bosso-Lefèvre, Y. Nadjar, A. Dumoulin, A. Triller, J.-L. Bessereau, L. Rondi-Reig, P. Isope, F. Selimi

## Abstract

Fine control of protein stoichiometry at synapses underlies brain function and plasticity. How proteostasis is controlled independently for each type of synaptic protein in a synapse-specific and activity-dependent manner remains unclear. Here we show that SUSD4, a complement-related transmembrane protein, binds the AMPA receptor subunit GLUA2 and controls its activity-dependent degradation. Several proteins with known roles in the regulation of AMPA receptor turnover, in particular ubiquitin ligases of the NEDD4 subfamily, are identified as SUSD4 binding partners. SUSD4 is expressed by many neuronal populations starting at the time of synapse formation. Loss-of-function of *Susd4* in the mouse prevents long-term depression at cerebellar synapses, and leads to impairment in motor coordination adaptation and learning. Our findings reveal that activity-dependent synaptic plasticity relies on a transmembrane CCP domain-containing protein that regulates the degradation of specific substrates. This mechanism potentially accounts for the role of SUSD4 mutations in neurodevelopmental diseases.

## Introduction

Proteostasis is at the core of many cellular processes and its dynamics needs to be finely regulated for each protein in each organelle. In neurons, additional challenges are imposed by their spatial complexity. In particular, during long-term synaptic plasticity, the proposed substrate for learning and memory (Collingridge et al., 2010; Nicoll, 2017), the number of neurotransmitter receptors needs to be regulated independently in a synapse-specific and activity-dependent manner. At excitatory synapses, the modification of AMPA receptor numbers is a highly dynamic process, involving regulation of receptor diffusion (Choquet and Triller, 2013; Penn et al., 2017), their insertion in the plasma membrane, anchoring at the postsynaptic density and endocytosis (Anggono and Huganir, 2012). After activity-dependent endocytosis, AMPA receptors are either recycled to the plasma membrane or targeted to the endolysosomal compartment for degradation (Ehlers, 2000; Lee et al., 2004; Park et al., 2004). The decision between these two fates, recycling or degradation, regulates the direction of synaptic plasticity. Recycling promotes long-term potentiation (LTP) and relies on many molecules, such as GRASP1, GRIP1, PICK1 and NSF (Anggono and Huganir, 2012). Targeting to the endolysosomal compartment and degradation promote long-term depression (LTD; Fernandez-Monreal et al., 2012; Kim et al., 2017; Matsuda et al., 2013), but the regulation of the targeting and degradation process remains poorly understood.

The Complement Control Protein domain (CCP), an evolutionarily conserved module also known as Sushi domain, was first characterized in proteins with role in immunity, in particular in the complement system. In the past few years, proteins with CCP domains have been increasingly recognized for their role at neuronal synapses. Acetylcholine receptor clustering is regulated by CCP domain-containing proteins in *Caenorhabditis elegans* (Gendrel et al., 2009) and in *Drosophila melanogaster* (Nakayama et al., 2016). In humans, mutations in the CCP domain-containing secreted protein SRPX2 are associated with epilepsy and speech dysfunction, and SRPX2 knockdown leads to decreased synapse number and vocalization in mice (Sia et al., 2013). Recently SRPX2 has been involved in the regulation of synapse elimination in the visual and somatosensory systems (Cong et al., 2020). Despite the increase in the diversity of CCP domain-containing proteins in evolution (11 CCP domain-containing in *C. elegans* and 56 in humans; <u>smart.embl.de</u>), the function of many CCP domain-containing proteins remains unknown.

The mammalian *SUSD4* gene codes for a transmembrane protein with four extracellular CCP domains (**Figure 1A**) and is highly expressed in the central nervous system (Holmquist et al., 2013). The *SUSD4* gene is located in a genomic region deleted in patients with the 1q41q42 syndrome that includes developmental delays and intellectual deficiency (ID; Rosenfeld et al., 2011). *SUSD4* is also amongst the 124 genes enriched in *de novo* missense mutations in a large cohort of individuals with Autism Spectrum Disorders (ASDs) or IDs (Coe et al., 2019). A copy number variation and several *de novo* mutations with a high CADD score, which indicates the deleteriousness of the mutations, have been described in the *SUSD4* gene in patients with ASDs ((Cuscó et al., 2009); denovo-db, Seattle, WA (<u>denovo-db.gs.washington.edu</u>) 10, 2019). The SUSD4 protein has been described to regulate complement system activation in erythrocytes by binding the C1Q globular domain (Holmquist et al., 2013). Interestingly, this domain is found in major synaptic regulators such as C1QA (Stevens et al., 2007), CBLNs (Matsuda et al., 2010; Uemura et al., 2010) and C1Q-like proteins (Bolliger et al., 2011; Kakegawa et al., 2015; Sigoillot et al., 2015). Altogether these studies point to a potential role of SUSD4 in synapse formation and/or function and in the etiology of neurodevelopmental disorders.

**Figure 1.**
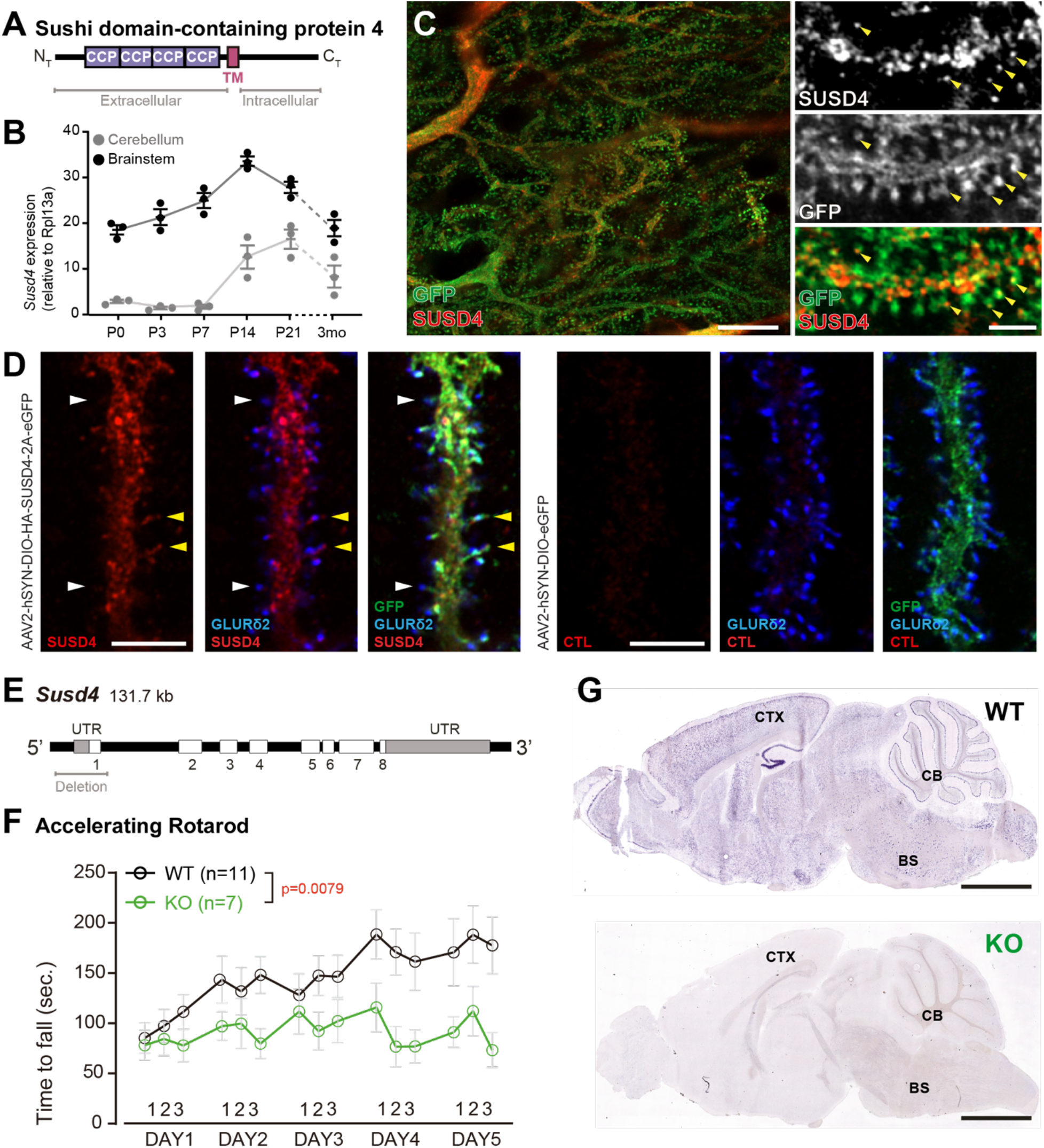
SUSD4 is necessary for motor coordination adaptation and learning. **(A)** Diagram of the protein SUSD4 showing its domain organization with four extracellular Complement Control Protein (CCP) domains, one transmembrane (TM) domain and a cytoplasmic domain (C_T_). **(B)** Quantitative RT-PCR shows an increase in *Susd4* mRNA expression (relative to the housekeeping gene *Rpl13a*) during postnatal development in the cerebellum and in the brainstem. Extracts were prepared from tissue samples of mice aged from 0 to 21 days (P0-21) and three months (3mo). Mean ± s.e.m. (n=3 independent experiments). **(C)** HA-tagged SUSD4 is found in dendrites (left panel, single plane) and in some of the distal dendritic spines (right panel, arrowheads, projection of a 1,95µm z-stack) in adult cerebellar Purkinje cells. Anti-HA and anti-GFP immunolabeling was performed on parasagittal cerebellar sections obtained from adult L7-Cre mice after stereotaxic injection of AAV particles driving the expression of HA-SUSD4 and soluble GFP. Scale bars: 10 µm (left panel) and 2µm (right panel). **(D)** Purkinje cells from primary mixed cerebellar culture of L7-Cre mice were transduced at 3 days in vitro (DIV3) with a HA-tagged SUSD4 expressing virus (AAV2-hSYN-DIO-HA-SUSD4-2A-eGFP) or with a control virus expressing GFP (AAV2-hSYN-DIO-eGFP), and immunostained in non-permeabilizing conditions at DIV17 for HA to localize surface SUSD4 (anti-HA, red), and in permeabilizing conditions to detect the green fluorescent protein (anti-GFP, green) and the endogenous GluD2 subunit (anti-GRID2, blue). Scale bar: 5 µm. **(E)** Genomic structure of the *Susd4* gene. White boxes represent exons. Exon 1 is deleted in the *Susd4* loss-of-function mouse model. See also **Figure S2**. **(F)** Motor coordination and learning is deficient in adult male *Susd4* ^-/-^ (KO) mice compared to age-matched *Susd4*^+/+^ (WT) littermates. Each mouse was tested three times per day during five consecutive days on an accelerating rotarod (4 to 40 r.p.m. in 10 minutes) and the time spent on the rotarod was measured. Mean ± s.e.m. (WT n=11 and KO n=7 mice, two-way ANOVA with repeated measures, Interaction (time and genotype): ** P=0.0079, F(14, 224) = 2.22; Time: **** P<0.0001, F(14, 224) = 3.469; Genotype: P=0.0553, F(1, 16) = 4.272). **(G)** *In situ* hybridization experiments were performed on brain sections from one month-old WT and *Susd4* KO mice to detect *Susd4* mRNA using a probe encompassing exons 2 to 5 (See also **Figure S2)**. *Susd4* expression was found in many regions of the brain in *Susd4*^+/+^ (WT) mice (see also **Figure S1**) including the cerebral cortex (CTX), the cerebellum (CB), and the brainstem (BS). No labeling was found in the brain of *Susd4* ^-/-^ (KO) mice. Scale bars: 500 µm.

Proper development and function of the cerebellar circuitry is central for motor coordination and adaptation, and a range of cognitive tasks (Badura et al., 2018; Hirai et al., 2005; Ichise et al., 2000; Lefort et al., 2019; Rochefort et al., 2011; Tsai et al., 2012). Cerebellar dysfunction is associated with several neurodevelopmental disorders including ASDs (Stoodley, 2016; Stoodley et al., 2018; Wang et al., 2014). In this circuit, cerebellar Purkinje cells (PCs) receive more than a hundred thousand parallel fiber (PF) synapses whose formation, maintenance and plasticity are essential for cerebellar-dependent learning (Gutierrez-Castellanos et al., 2017; Hirai et al., 2005; Ito, 2006; Kashiwabuchi et al., 1995). Postsynaptic LTD was first described at synapses between PFs and cerebellar PCs (Gao et al., 2012; Hirano, 2018; Ito, 2001; Ito and Kano, 1982), where it can be induced by conjunctive stimulation of PFs with the other excitatory input received by PCs, the climbing fiber (CF; Coesmans et al., 2004; Ito, 2001; Suvrathan et al., 2016). The function of members of the C1Q family, such as CBLN1 and C1QL1, is essential for excitatory synapse formation and LTD in cerebellar PCs (Hirai et al., 2005; Kakegawa et al., 2015; Matsuda et al., 2010; Sigoillot et al., 2015; Uemura et al., 2010), suggesting that proteins such as SUSD4, that interact with the C1Q globular domain, could regulate these processes.

Gene expression studies from our laboratory revealed that *Susd4* is highly expressed in the olivocerebellar system of the mouse. In order to uncover the potential link between SUSD4 and neurodevelopmental disorders and the involvement of the olivocerebellar circuit in such diseases, we sought to identify the role of SUSD4 in brain development and function, by analyzing the phenotype of a *Susd4* loss-of-function mouse model in the cerebellum. Here we show that knockout of the *Susd4* gene leads to misregulation of synaptic plasticity in cerebellar PCs, deficits in motor coordination adaptation and learning as well as an impairment in the control of activity-dependent degradation of GLUA2 AMPA receptor subunits. Using affinity-purification of synaptosome preparations followed by proteomics analysis, we found that the SUSD4 protein binds proteins that are involved in the regulation of several parameters controlling AMPA receptor turnover. In particular, SUSD4 directly interacts with E3 ubiquitin ligases of the NEDD4 family, which are known to regulate ubiquitination and degradation of their substrates. Finally, our results suggest that SUSD4 promotes degradation of GLUA2 subunits over recycling at synapses, thereby contributing to the decision between potentiation and depression. Given the domain structure of SUSD4, this new regulatory mechanism could bring spatio-temporal specificity to the degradation machinery in neurons, allowing proper synaptic plasticity and learning.

## Results

### *Susd4* is broadly expressed in neurons during postnatal development

Given the potential synaptic role for SUSD4, its pattern of expression should correlate with the timing of synapse formation and/or maturation during postnatal development. *In situ* hybridization experiments using mouse brain sections showed high expression of *Susd4* mRNA in neurons in many regions of the central nervous system, including the cerebral cortex, the hippocampus, the cerebellum and the brainstem (**Figure 1B** and **S1**). *Susd4* expression was already detected as early as postnatal day 0 (P0) in some regions, but increased with brain maturation (**Figure S1**). In the cerebellum, a structure where the developmental sequence leading to circuit formation and maturation is well described (Sotelo, 2004), quantitative RT-PCR showed that *Susd4* mRNA levels start increasing at P7 and by P21 reach about 15 times the levels detected at birth (**Figure 1B**). At P7, a major increase in synaptogenesis is observed in the cerebellum. At this stage, hundreds of thousands of PF excitatory synapses form on the distal dendritic spines of each PC, and a single CF arising from an inferior olivary neuron translocates and forms about 300 excitatory synapses on proximal PC dendrites (Leto et al., 2016). In the brainstem, where cell bodies of inferior olivary neurons are located, the increase in *Susd4* mRNA expression occurs earlier, already by P3, and reaches a peak by P14 (**Figure 1B**). Similarly to the cerebellum, this pattern of *Susd4* expression parallels the rate of synaptogenesis that increases during the first postnatal week in the inferior olive (Gotow and Sotelo, 1987). To identify the subcellular localization of the SUSD4 protein and because of the lack of suitable antibodies for immunolabeling, viral particles enabling CRE-dependent coexpression of HA-tagged SUSD4 and GFP in neurons were injected in the cerebellum of adult mice expressing the CRE recombinase specifically in cerebellar PCs. Immunofluorescent labeling against the HA tag demonstrated the localization of HA-SUSD4 in dendrites and in some of the numerous dendritic spines present on the surface of distal dendrites (**Figure 1C**). These spines are the postsynaptic compartments of PF synapses in PCs. Immunofluorescence analysis of transduced cultured PCs further showed that HA-tagged SUSD4 could be immunolabeled in non-permeabilizing conditions and located at the surface of dendrites and spines **(Figure 1D)**. Double labeling with the postsynaptic marker GLURδ2 (GRID2) further showed partial colocalization at the surface of some, but not all, spines. Therefore, the timing of *Susd4* mRNA expression during postnatal development and the subcellular localization of the SUSD4 protein in cerebellar PCs are in agreement with a potential role for SUSD4 in excitatory synapse formation and/or function.

### *Susd4* loss-of-function leads to deficits in motor coordination and learning

To determine the synaptic function of SUSD4, we analyzed the phenotype of *Susd4*^*-/-*^ constitutive knockout (KO) mice with a deletion of exon 1 (**Figure 1E, 1G** and **S2**). RT-PCR using primers encompassing the last exons and the 3’UTR show the complete absence of *Susd4* mRNA in the brain of these *Susd4* KO mice (**Figure S2**). No obvious alterations of mouse development and behavior were detected in those mutants, an observation that was confirmed by assessment of their physical characteristics (weight, piloerection), basic behavioral abilities such as sensorimotor reflexes (whisker responses, eye blinking) and motor responses (open field locomotion; cf. **Table S1**). Because of the high expression of *Susd4* in the olivocerebellar system (**Figures 1G** and **S1**), we further assessed the behavior of *Susd4* KO mice for two abilities well known to depend on normal function of this network, motor coordination and motor learning (Kayakabe et al., 2014; Lalonde and Strazielle, 2001; Rondi-Reig et al., 1997). Using a footprint test, a slightly larger print separation of the front and hind paws in the *Susd4* KO mice was detected but no differences in the stride length and stance width were found (**Figure S3**). In the accelerated rotarod assay, a classical test of motor adaptation and learning (Buitrago et al., 2004), the mice were tested three times per day at one hour interval during five consecutive days. The *Susd4* KO mice performed as well as the *Susd4*^*+/+*^ (WT) littermate controls on the first trial (**Figure 1F, day 1, trial 1**). This indicates that there is no deficit in their balance function, despite the slight change in fine motor coordination found in the footprint test. However, while the control mice improved their performance as early as the third trial on the first day, and further improved with several days of training, no learning could be observed for the *Susd4* KO mice either during the first day, or in the following days (**Figure 1F)**. These results show that *Susd4* loss-of-function leads to impaired motor coordination and learning in adult mice.

### *Susd4* loss-of-function prevents long-term depression (LTD) at cerebellar parallel fiber/Purkinje cell synapses

Motor coordination and learning are deficient when cerebellar development is impaired (Hirai et al., 2005; Ichise et al., 2000; Tsai et al., 2012; Zuo et al., 1997). No deficits in the global cytoarchitecture of the cerebellum and morphology of PCs were found in *Susd4* KO mice (**Figure S4**). Using high density microelectrode array, we assessed the spontaneous activity of PCs in acute cerebellar slices from *Susd4* KO mice, and compared to *Susd4* WT mice (**Figure S5**). No differences were detected in either the mean spiking frequency, the coefficient of variation of interspike intervals (CV) and the intrinsic variability of spike trains (CV2, Holt and Douglas, 1996) indicating that the firing properties of PCs are not affected by *Susd4* loss-of-function.

Co-immunolabeling of PF presynaptic boutons using an anti-VGLUT1 antibody and PCs using an anti-calbindin antibody in cerebellar sections from juvenile WT mice revealed an extremely dense staining in the molecular layer corresponding to the highly numerous PFs contacting PC distal dendritic spines (**Figure 2A**). The labeling pattern appeared to be similar in *Susd4* KO. High-resolution microscopy and quantitative analysis confirmed that there are no significant changes in the mean density and volume of VGLUT1 clusters following *Susd4* loss-of-function (**Figure 2A**). Electric stimulation of increasing intensity in the molecular layer allows the progressive recruitment of PFs (Konnerth et al., 1990), and can be used to assess the number of synapses and basic PF/PC transmission using whole-cell patch-clamp recordings of PCs on acute cerebellar slices (**Figure 2B**). No difference was observed in the amplitude and the kinetics of the responses to PF stimulation in PCs from *Susd4* KO and control littermate mice (**Figure 2C** and **Figure S6**). Furthermore, the probability of vesicular release in the presynaptic PF boutons, as assessed by measurements of paired pulse facilitation (Atluri and Regehr, 1996; Konnerth et al., 1990; Valera et al., 2012), was not changed at PF/PC synapses (**Figure 2C**). Finally, no differences in the frequency and amplitude of PF/PC evoked quantal events were detected (**Figure S6**). Thus, in accordance with the morphological analysis, *Susd4* invalidation has no major effect on the number and basal transmission of PF/PC synapses in the mouse.

**Figure 2.**
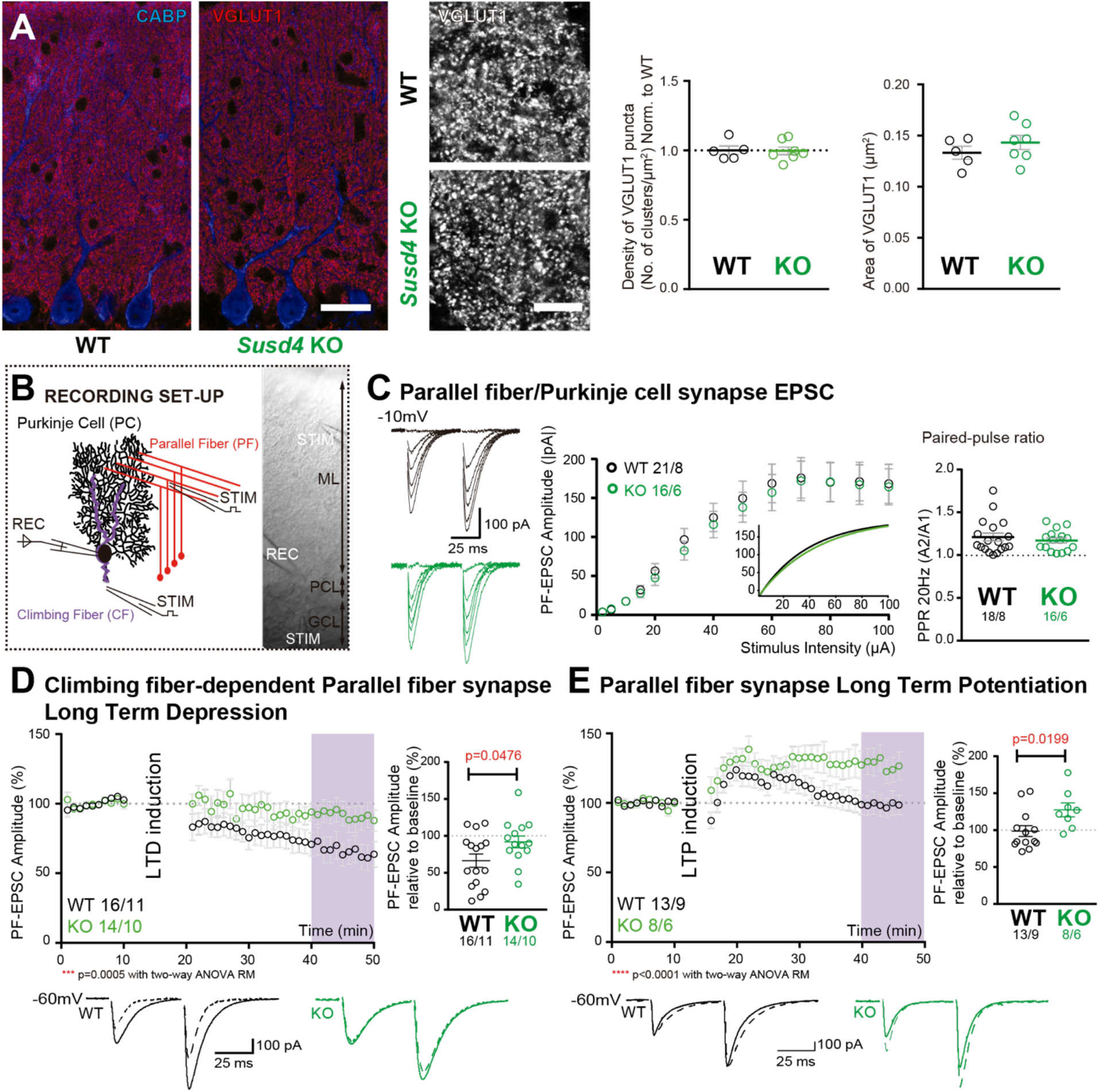
*Susd4* loss-of-function leads to deficient long-term depression and facilitated long-term potentiation of parallel fiber/Purkinje cell synapses. **(A)** Quantitative analysis of the morphology of parallel fiber presynaptic boutons immunolabeled by an anti-VGLUT1 antibody (red) in Purkinje cells (anti-CABP, blue). Quantifications of the density and the area of the VGLUT1 clusters did not reveal any difference between *Susd4* KO and *Susd4* WT mice. Mean ± s.e.m. (WT n=5 and KO n=7 mice; VGLUT1 clusters density: Mann-Whitney test, P>0.9999; area VGLUT1 clusters: Unpaired Student t-test, P=0.3089). Scale bars: 30 µm (left) and 10 µm (right). **(B)** Diagram of the setup for patch-clamp recordings (REC) of Purkinje cells in 300 µm-thick parasagittal cerebellar slices. Parallel fiber and climbing fiber responses were elicited by electrical stimulation (STIM). ML: molecular layer; PCL: Purkinje cell layer; GCL: granule cell layer. **(C)** Input-output curve of the parallel fiber/Purkinje cell transmission. The amplitude of the elicited EPSCs increases with the intensity of the stimulus and is not significantly different between *Susd4* KO and WT littermates. The fitted curves for each genotype are presented in the inset. Representative sample traces are presented. Mean ± s.e.m. (WT n=18 cells from 8 mice and KO n=16 cells from 6 mice; Kolmogorov-Smirnov test, P=0.8793). Short-term plasticity of parallel fiber/Purkinje cell synapses is not affected by *Susd4* loss-of-function. Parallel fibers were stimulated twice at 50 ms interval and the paired-pulse ratio (PPR) was calculated by dividing the amplitude of the second peak by the amplitude of the first peak. Mean ± s.e.m. (WT n=21 cells from 8 mice and KO n=16 cells from 6 mice; Mann-Whitney test, P=0.9052). **(D)** Climbing fiber-dependent parallel fiber/Purkinje cell synapse long-term depression (LTD) is impaired in the absence of *Susd4* expression. LTD was induced by pairing stimulations of parallel fibers and climbing fibers at 100 milliseconds interval during 10 minutes at 0.5 Hz (see also **Figure S6**). The amplitude of the PF EPSC was measured using two consecutive PF stimulation at 50 milliseconds interval. Representative sample traces are presented. Right: EPSC amplitudes from the last 10 minutes (purple) of recordings were used to calculate the LTD ratio relative to baseline. Mean ± s.e.m. (WT n=16 cells from 11 mice and KO n=14 cells from 10 mice; Two-tailed Wilcoxon Signed Rank Test with null hypothesis of 100: WT **p=0.0063; KO p=0.2676; Mann-Whitney test, WT *vs* KO *p=0.0476). **(E)** Loss-of-function of *Susd4* facilitates parallel fiber/Purkinje cell synapse long-term potentiation (LTP). Tetanic stimulation of only parallel fibers at 0.3 Hz for 100 times (see also **Figure S6**) induced LTP in *Susd4* KO Purkinje cells while inducing only a transient increase in parallel fiber transmission in WT Purkinje cells. Representative sample traces are presented. Right: EPSC amplitudes from the last 7 minutes (purple) were used to calculate the LTP ratio relative to baseline. Mean ± s.e.m. (WT n=13 cells from 9 mice and KO n=8 cells from 6 mice; Two-tailed Wilcoxon Signed Rank Test with null hypothesis of 100: WT p=0.5879; KO *p=0.0234; Mann-Whitney test, WT *vs* KO: *p=0.0199).

Long-term synaptic plasticity of PF/PC synapses is involved in proper motor coordination and adaptation learning (Gutierrez-Castellanos et al., 2017; Hirano, 2018; Kakegawa et al., 2018). We first assessed LTD in PF/PC synapses using conjunctive stimulation of PFs and CFs and whole-cell patch-clamp recordings of PCs in acute cerebellar slices from juvenile mice. The LTD induction protocol produced a 42% average decrease in the amplitude of PF excitatory postsynaptic currents (EPSCs) in PCs from WT mice while the paired pulse facilitation ratio was not changed during the course of our recordings (**Figure 2D** and **Figure S6**). In *Susd4* KO PCs, the same LTD induction protocol did not induce any significant change in PF EPSCs during the 30 minutes recording period, showing that LTD induction and maintenance are greatly impaired in the absence of SUSD4 (**Figure 2D**). LTP can be induced by high frequency stimulation of PFs only, and is also involved in cerebellar dependent-learning (Binda et al., 2016; Gutierrez-Castellanos et al., 2017). In control mice, tetanic stimulation during 5 minutes induced a transient increase in transmission of about 20% and the amplitude of the response returned to baseline after only 15 minutes (**Figure 2E** and **Figure S6**). However, in the case of *Susd4* KO PCs, the same protocol induced a 27% increase in transmission that was maintained after 35 minutes (**Figure 2E**), indicating a facilitation of LTP of PF/PC synapses in the absence of *Susd4* expression.

Lack of LTD of PF/PC synapses could arise from deficient CF/PC transmission. To test this possibility, we first crossed the *Susd4* KO mice with the *Htr5b*-GFP BAC transgenic line (http://gensat.org/MMRRC_report.jsp?founder_id=17735) expressing soluble GFP specifically in inferior olivary neurons in the olivocerebellar system to visualize CFs. We found that CFs had a normal morphology and translocated along the proximal dendrites of their PC target in *Susd4* KO mice (**Figure 3A**). We then assessed whether developmental elimination of supernumerary CFs was affected by *Susd4* invalidation using whole-cell patch-clamp recordings of PCs on cerebellar acute slices (Crepel et al., 1976; Hashimoto and Kano, 2003). No difference was found in the percentage of remaining multiply-innervated PCs in the absence of *Susd4* (**Figure 3B**). We next used VGLUT2 immunostaining to label CF presynaptic boutons and analyze their morphology using high resolution confocal microscopy and quantitative image analysis. VGLUT2 immunostaining revealed the typical CF innervation territory on PC proximal dendrites, extending up to about 80% of the molecular layer height both in control *Susd4* WT and in *Susd4* KO mice (**Figure 3C**). Furthermore, the number and density of VGLUT2 clusters were not significantly different between *Susd4* WT and *Susd4* KO mice. To test whether the lack of CF-dependent PF LTD was due to deficient CF transmission, we used whole-cell patch-clamp recordings of PCs in acute cerebellar slices. Contrary to what could have been expected, the typical all-or-none CF evoked EPSC was detected in PCs from *Susd4* KO mice with increased amplitude when compared to WT PCs (**Figure 3D**) while no differences in CF-EPSC kinetics were found (**Figure S7**). Therefore, the lack of CF-dependent PF/PC synapse LTD in *Susd4* KO mice is not due to impaired CF/PC synapse formation or transmission. Measurements of evoked quantal events revealed an increase in the amplitude of the quantal EPSCs at CF/PC synapses from juvenile mice (**Figures 3E** and **S7**). Paired-pulse facilitation and depression at PF/PC and CF/PC synapses, respectively, are similar between *Susd4* KO and control mice, both in basal conditions and during plasticity recordings (**Figures 2C, 3D, S6D** and **S6F**) suggesting strongly that the changes in PF/PC synaptic plasticity and in CF/PC transmission in *Susd4* KO PCs have a postsynaptic origin. Overall our results show that *Susd4* loss-of-function in mice leads to a highly specific phenotype characterized by misregulation of postsynaptic plasticity in the absence of defects in synaptogenesis and in basal transmission in cerebellar PCs.

**Figure 3.**
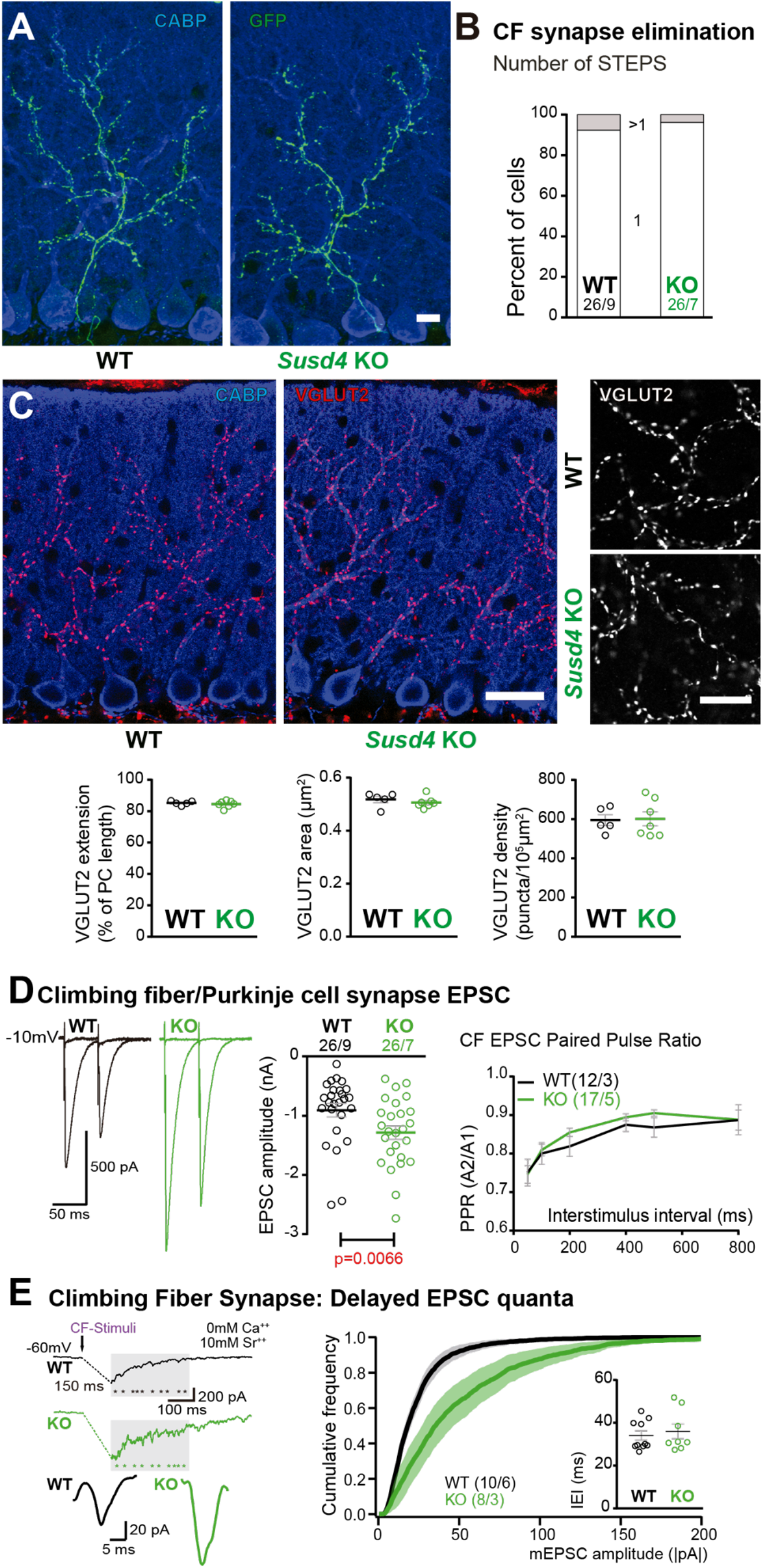
Transmission at the Climbing fiber/Purkinje cell synapses is increased in *Susd4* knockout mice. **(A)** Left: Climbing fibers were visualized in *Susd4* WT and KO mice crossed with Htr5b-eGFP reporter mice expressing the green fluorescent protein specifically in inferior olivary neurons. Anti-GFP and anti-CABP (to visualize Purkinje cells) immunofluorescence was performed on parasagittal sections of P30 mice, and showed no qualitative differences in the absence of *Susd4* expression. Scale bar: 10 µm. **(B)** Patch-clamp recordings of Purkinje cells showed a similar percentage of mono-(1 climbing fiber) and multi-innervation (>1 climbing fibers) of Purkinje cells in P30 *Susd4* KO and WT mice, as measured by the number of steps elicited by electrical stimulation of the climbing fibers. (WT n=26 cells from 9 mice and KO n=26 cells from 7 mice; Chi-square test, P=0.5520). **(C)** Climbing fiber presynaptic boutons were immunostained with an anti-VGLUT2 antibody in cerebellar sections from P30 WT and *Susd4* KO mice. The extension of the climbing fiber territory was calculated by measuring the extent of the VGLUT2 (red) labeling relative to the height of the Purkinje cell dendritic tree (immunostained using an anti-CABP antibody, blue). Quantification of the mean density of VGLUT2 puncta and their mean area showed no differences between *Susd4* KO mice and their control littermates. Mean ± s.e.m. (WT n=5 and KO n=7 mice; VGLUT2 extension: Mann-Whitney test, P=0.6389; VGLUT2 area: Unpaired Student t-test, p=0.4311; VGLUT2 density: Unpaired Student t-test, p=0.8925). Scale bars: 30 µm (left) and 10 µm (right). **(D)** Short-term synaptic plasticity of climbing fiber/Purkinje cell synapses was elicited by two consecutive stimulations at various intervals. The amplitude of the climbing fiber elicited EPSC was increased in *Susd4* KO mice compared to WT littermates. (WT n=26 cells, 9 mice and KO n=26 cells, 7 mice, Mann-Whitney test, ** P=0.0066). No difference in the paired pulse ratios (PPR) was detected at any interval between *Susd4* KO mice and WT mice. Representative sample traces are presented. See also **Figure S7**. Mean ± s.e.m. (WT n=12 cells from 3 mice and KO n=17 cells from 5 mice; Kolmogorov-Smirnov test, P=0.4740). **(E)** Delayed CF-EPSC quanta were evoked by CF stimulation in the presence of Sr^++^ instead of Ca^++^ to induce desynchronization of fusion events. Representative sample traces are presented. The cumulative probability for the amplitude of the events together with the individual amplitude values for each event show an increased amplitude associated with *Susd4* loss-of-function. The individual frequency values for each cell (measured as interevent interval, IEI) present no differences between the genotypes. See also **Figure S7**. Mean ± s.e.m. (WT n=10 cells from 6 mice and KO n=8 cells from 3 mice; Amplitude: Kolmogorov-Smirnov distribution test, *** P<0.0001; Frequency: Mann Whitney test, P=0.6334).

### *Susd4* loss-of-function leads to deficient activity-dependent degradation of GLUA2

What are the mechanisms that allow regulation of long-term synaptic plasticity by SUSD4? The lack of LTD at PF/PC synapses and our analysis of evoked quantal events suggested the involvement of SUSD4 in the regulation of postsynaptic receptor numbers. GLUA2 subunits are present in most AMPA receptor channels in PC excitatory synapses (Masugi-Tokita et al., 2007; Zhao et al., 1998). To assess whether *Susd4* loss-of-function leads to misregulation of the GLUA2 subunits at PC excitatory synapses, we first performed co-immunolabeling experiments using an anti-GLUA2 antibody and an anti-VGLUT2 antibody on cerebellar sections followed by high-resolution microscopy. Several GLUA2 clusters of varying sizes were detected in close association with each VGLUT2 presynaptic cluster corresponding to a single CF release site, while very small and dense GLUA2 clusters were found in the rest of the molecular layer which mostly correspond to GLUA2 clusters at the PF/PC synapses (**Figure 4A**). No obvious change in GLUA2 distribution in the molecular layer in *Susd4* KO mice was found when compared to controls, in accordance with normal basal transmission in PF/PC synapses (**Figure 2C**). Quantitative analysis of the GLUA2 clusters associated with VGLUT2 labelled CF presynaptic boutons did not reveal a significant change in the total mean intensity of GLUA2 clusters per CF presynaptic bouton (**Figure 4A**). However, the proportion of CF presynaptic boutons with no GLUA2 cluster was smaller in juvenile *Susd4* KO mice than in WT mice (**Figure 4A**). This decrease partially explains the increase in the amplitude of quantal EPSCs and CF transmission (**Figure 3E**).

**Figure 4.**
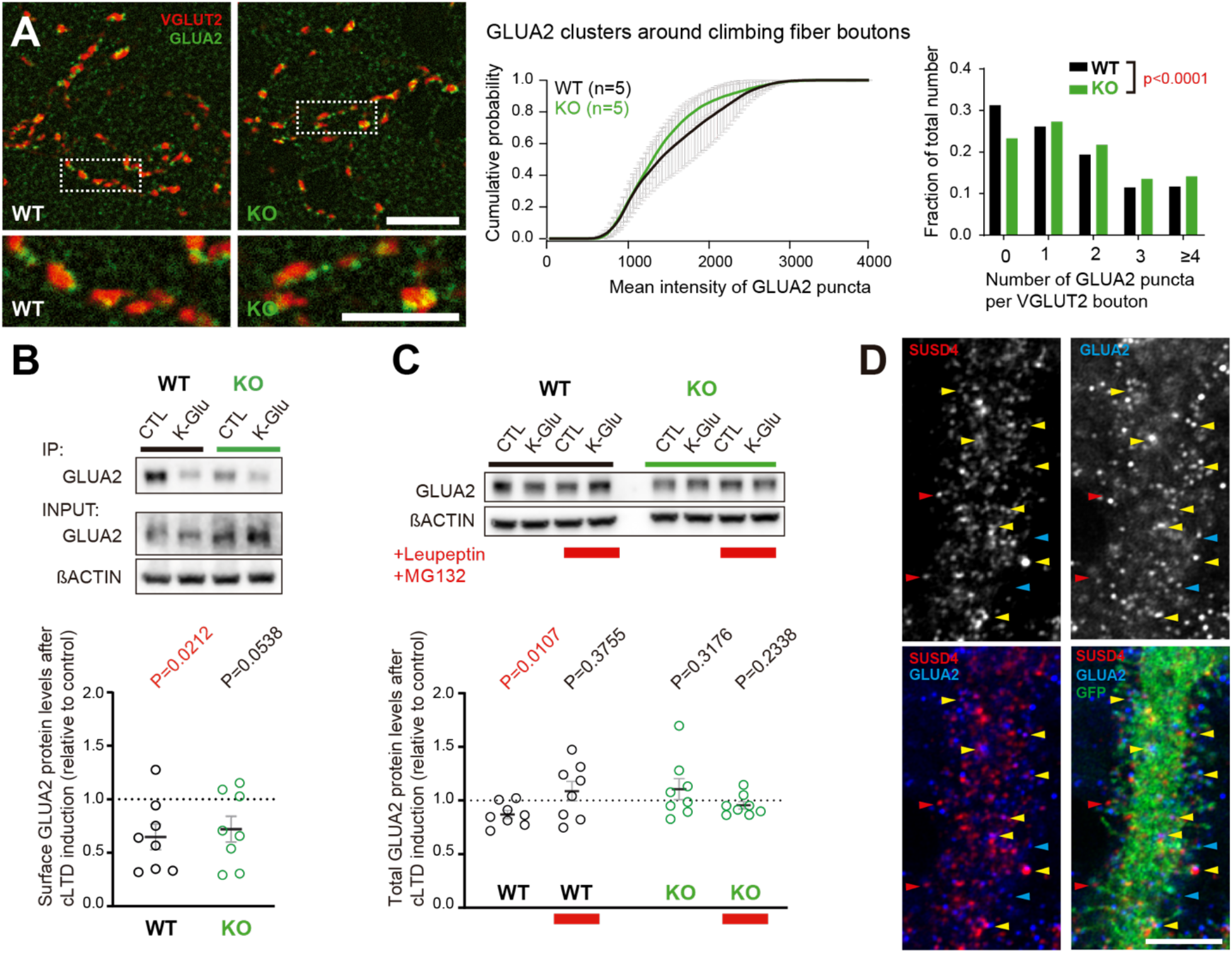
Loss of SUSD4 leads to misregulation of the AMPA receptor subunit GLUA2. **(A)** The number of GLUA2 clusters (anti-GLUA2 immunolabeling, green) per climbing fiber presynaptic bouton (anti-VGLUT2 immunolabeling, red) and their intensity were quantified in cerebellar sections of juvenile *Susd4*^*-/-*^ KO mice and *Susd4*^*+/+*^ WT littermates. Cumulative plot for the mean GLUA2 intensity per VGLUT2 bouton shows no significant change between WT and KO. The distribution of the VGLUT2 boutons according to the number of associated GLUA2 clusters is significantly different between WT and KO. Mean ± s.e.m. (WT n= 5 and KO n= 5 mice; Intensity: Kolmogorov-Smirnov test, P=0.5009; Distribution: Chi-square contingency test, **** P<0.0001). Scale bars: 30 µm (top) and 15 µm (bottom). **(B)** Activity-dependent changes in surface localization of GLUA2 was studied in cerebellar acute slices from *Susd4* KO mice and control *Susd4* WT littermates using a chemical LTD protocol (cLTD; K-Glu: K^+^ 50mM and glutamate 10µM for 5 minutes followed by 30 minutes of recovery). Surface biotinylation of GLUA2 subunits was performed followed by affinity-purification of biotinylated GLUA2 subunits and anti-GLUA2 immunoblot analysis. The fraction of biotinylated GLUA2 was obtained by measuring the levels of biotinylated GLUA2 in affinity-purified samples and total GLUA2 normalized to beta-actin in input samples for each condition. The ratios between the fraction of biotinylated GLUA2 after cLTD and control conditions are represented. Mean ± s.e.m. (n=8 independent experiments; Two-tailed Student’s one sample t-test was performed on the ratios with a null hypothesis of 1, P_WT_ = 0.0212 and P_KO_ = 0.0538). **(C)** Activity-dependent degradation of GLUA2 was assessed in cerebellar acute slices from *Susd4* KO and control mice after induction of chemical LTD (cLTD; K-Glu: K^+^ 50mM and glutamate 10µM for 5 minutes followed by 30 minutes of recovery). This degradation was absent when slices were incubated with 100µg/mL leupeptin and with 50µM MG132 (to inhibit lysosomal and proteasome degradation, respectively), or when slices were obtained from *Susd4* KO mice. Band intensities of GLUA2 were normalized to β-ACTIN. The ratios between levels with cLTD induction (K-Glu) and without cLTD induction (CTL) are represented. See also **Figure S8**. Mean ± s.e.m. (n=8 independent experiments; Two-tailed Student’s one sample t-test was performed on the ratios with a null hypothesis of 1, P_WT_ = 0.0107, P_WT+Leu/MG132_ = 0.3755, P_KO_ = 0.3176 and P_KO+Leu/MG132_ = 0.2338). **(D)** Purkinje cells from primary cerebellar cultures of L7-Cre mice were transduced at 3 days in vitro (DIV3) with a virus driving expression of HA-tagged SUSD4 (AAV2-hSYN-DIO-HA-SUSD4-2A-eGFP) and immunolabeled at DIV17 in non-permeabilizing conditions to localize surface SUSD4 (anti-HA, red) and surface GLUA2 subunits (anti-GluA2, blue). Direct green fluorescent protein is shown (GFP, green). Examples of spines containing either SUSD4 alone, GLUA2 alone or both are shown using red, blue and yellow arrowheads, respectively (maximum projection of a 1.8 µm z-stack). Scale bar: 5 µm.

In cerebellar PCs, regulation of the GLUA2 subunits at synapses and of their trafficking is essential for PF LTD (Chung et al., 2003; Xia et al., 2000). To test whether activity-dependent surface localization of GLUA2-containing AMPA receptors is affected by loss of *Susd4*, we set up a biochemical assay in which we induced chemical LTD (cLTD) in acute cerebellar slices (Kim et al., 2017) and performed surface biotinylation of GLUA2 subunits followed by immunoblot quantification. As expected, our results showed a 35% mean reduction of surface GLUA2 receptors after cLTD in slices from WT mice (**Figure 4B**). In slices from *Susd4* KO mice, a similar mean reduction of surface GLUA2 receptors was detected after cLTD (28%), suggesting that SUSD4 does not affect the machinery controlling endocytosis of GLUA2 subunits. Another parameter that needs to be controlled for proper LTD in PCs is the total number of AMPA receptors in the recycling pool and the targeting of AMPA receptors to late endosomes and lysosomes (Kim et al., 2017). Lack of LTD and facilitation of LTP in *Susd4* KO mice (**Figures 2D** and **2E**) suggest that GLUA2 activity-dependent targeting to the endolysosomal compartment and its degradation is affected by *Susd4* loss-of-function. Using our cLTD assay in cerebellar slices, we measured the total GLUA2 levels either in control conditions or in presence of inhibitors of the proteasome (MG132) and of lysosomal degradation (leupeptin). The comparison of the GLUA2 levels in the presence of both inhibitors and in control conditions allowed us to estimate the GLUA2 degraded pool, regardless of the mechanism behind this degradation. On average, total GLUA2 levels were not significantly different between *Susd4* WT and *Susd4* KO cerebellar slices in basal conditions (**Figure S8**), in accordance with our morphological and electrophysiological analysis of PF/PC synapses (**Figures 2A** and **2C**). In slices from WT mice, chemical induction of LTD induced a significant reduction of 13% in total GLUA2 protein levels (**Figure 4C**). This reduction was prevented by incubation with the mixture of degradation inhibitors, MG132 and leupeptin, showing that it corresponds to the pool of GLUA2 degraded in an activity-dependent manner (**Figure 4C**). In slices from *Susd4* KO mice, this activity-dependent degradation of GLUA2 was completely absent. Additionally, the chemical induction of LTD had no effect on the total protein levels of GLURδ2, another synaptic receptor highly present at PF/PC postsynaptic densities, either in slices from WT or from *Susd4* KO mice (**Figure S8**). Thus, SUSD4 specifically controls the activity-dependent degradation of GLUA2-containing AMPA receptors during LTD.

Finally, in order to assess the potential colocalization of SUSD4 and GLUA2 in neurons, we used a Cre-dependent AAV construct to express HA-tagged SUSD4 in cultured PCs (**Figure 4D**) and performed immunolabeling of surface GLUA2 subunits. Clusters of HA-tagged SUSD4 partially colocalize with GLUA2 clusters at the surface of some dendritic spines (yellow arrowheads, **Figure 4D**). Partial colocalization of GLUA2 and SUSD4 in neurons was also confirmed in transfection experiments in hippocampal neurons (**Figure S9**). Altogether, these results suggest that SUSD4 could regulate activity-dependent degradation of GLUA2-containing AMPA receptors through a direct interaction.

### SUSD4 could regulate the number of AMPA receptor at synapses via multiple molecular interactions

To better understand how SUSD4 might regulate the number of GLUA2-containing AMPA receptors at synapses, we searched for SUSD4 molecular partners by affinity-purification of cerebellar synaptosome extracts using GFP-tagged SUSD4 as a bait (**Figure 5A**). Interacting partners were identified by proteomic analysis using liquid chromatography with tandem mass spectrometry (LC-MS/MS; Savas et al., 2014). 28 candidates were identified including proteins with known function in the regulation of AMPA receptor turnover (**Figure 5E**). Several candidates were functionally linked to ubiquitin ligase activity by gene ontology term analysis (**Figure 5A** and **Table 1**). In particular, five members of the NEDD4 subfamily of HECT E3 ubiquitin ligases were found as potential interacting partners, three of them (*Nedd4l, Wwp1* and *Itch*) exhibiting the highest enrichment factors amongst the 28 candidates. Ubiquitination is a post-translational modification essential for the regulation of protein turnover and trafficking in cells (Tai and Schuman, 2008). A survey of the expression of HECT-ubiquitin ligases shows that different members of the NEDD4 subfamily are broadly expressed in the mouse brain, however with only partially overlapping patterns (**Figure S9**, http://mouse.brain-map.org, Allen Brain Atlas). *Nedd4* and *Wwp1* are the most broadly expressed, including in neurons that also express *Susd4*, such as hippocampal neurons, inferior olivary neurons in the brainstem and cerebellar PCs. SUSD4 interaction with NEDD4 ubiquitin ligases could thus participate in the control of synaptic plasticity and of GLUA2 AMPA receptor subunit degradation. Immunoblot analysis of affinity-purified synaptosome extracts confirmed the interaction of SUSD4 with NEDD4, ITCH and WWP1 (**Figure 5B**). Removal of the intracellular domain of SUSD4 (SUSD4*Δ*C_T_ mutant) prevented this interaction demonstrating the specificity of SUSD4 binding to NEDD4 ubiquitin ligases (**Figure 5B**).

**Figure 5.**
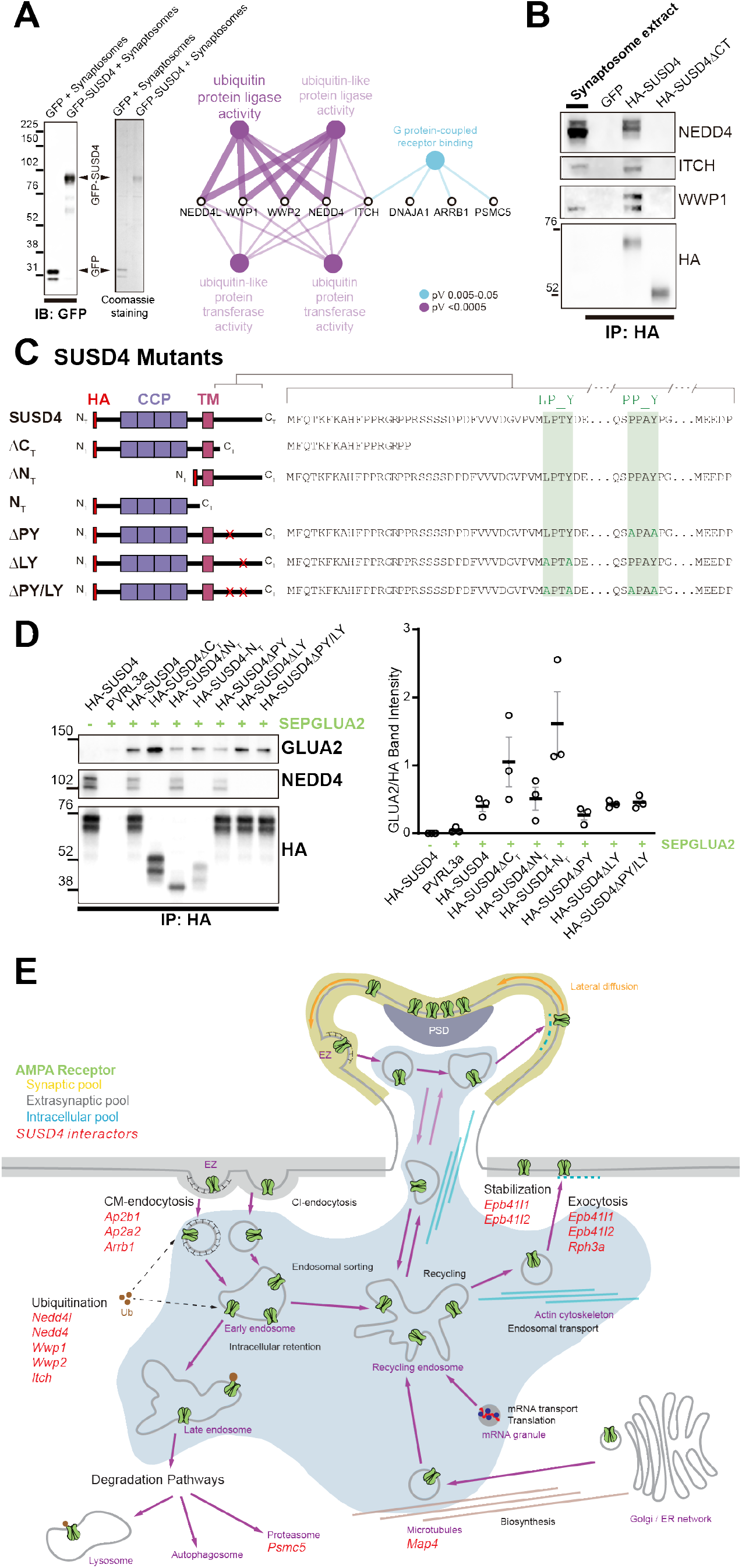
SUSD4 could regulate AMPA receptor degradation via multiple molecular interactions. **(A)** Mass spectrometry identification of SUSD4 interactors. Left: Affinity-purification from cerebellar synaptosomes was performed using either GFP-SUSD4 as a bait or GFP as a control. Proteins were then resolved using SDS-PAGE followed by immunoblot for anti-GFP and coomassie staining of proteins. Right: Gene Ontology (GO) enrichment analysis network (Molecular Function category) of the 28 candidate proteins (Cytoscape plugin ClueGO) identified after affinity-purification of cerebellar synaptosomes using GFP-SUSD4 as a bait followed by LC MS/MS. The Ubiquitin ligase activity term is significantly enriched in particular due to the identification of several members of the NEDD4 family of HECT-ubiquitin ligase. See also **Table 1**. (n=3 independent experiments). **(B)** Immunoblot confirmation of SUSD4 interaction with NEDD4 ubiquitin ligases. Affinity-purification from cerebellar synaptosomes was performed using either full length HA-SUSD4, HA-SUSD4ΔC_T_ or GFP as a bait. Proteins were then resolved using SDS-PAGE followed by immunoblot for NEDD4, ITCH, WWP1 or HA-SUSD4 (anti-HA). Full-length SUSD4 (HA-tagged, HA-SUSD4) interacts with all three members of the NEDD4 family. This interaction is lost when the C-terminal tail of SUSD4 is deleted (HA-SUSD4ΔC_T_) or when GFP is used instead of SUSD4 as a control. **(C)** Schematic representation of HA-tagged SUSD4 and different mutant constructs: SUSD4ΔC_T_ (lacking the cytoplasmic tail), SUSD4ΔN_T_ (lacking the extracellular domain), SUSD4N_T_ (lacking the transmembrane and intracellular domains), SUSD4ΔPY (point mutation of the PPxY site), SUSD4ΔLY (point mutation of the LPxY) and SUSD4ΔPY/LY (double mutant at both PPxY and LPxY). **(D)** SUSD4 interaction with GLUA2 and NEDD4 was assessed by co-immunoprecipitation using HEK293 cells transfected with SEP-GLUA2 together with PVRL3α as a control or one of the HA-SUSD4 constructs. Affinity-purification was performed with an anti-HA antibody and extracts were probed for co-immunoprecipitation of GLUA2 (with an anti-GLUA2 antibody) and of the HECT ubiquitin ligase NEDD4 (anti-NEDD4 antibody). The level of GLUA2 co-immunoprecipitated with each SUSD4 construct was quantified by performing the ratio of GLUA2 band intensity over the HA band intensity. N=3 independent experiments. **(E)** Potential interactors of SUSD4 control several parameters of AMPA receptor turnover. Three different pools of AMPA receptors are found in dendrites and spines: synaptic, extrasynaptic and intracellular. AMPA receptors are synthetized and delivered close to the synaptic spine to reach the synaptic surface. At the surface, AMPA receptors can move laterally (lateral diffusion) or vertically by endocytosis and exocytosis. Endocytosis can be mediated by clathrin (CM-endocytosis) or be clathrin-independent (CI-endocytosis). CM-endocytosis is often related to activity-dependent processes. After endocytosis, AMPA receptors can choose between two different pathways from the early endosomes, one for recycling and the other for degradation. Potential molecular partners of SUSD4 identified by our proteomics analysis could regulate AMPA receptor turnover at several levels of this cycle (in red).

One possibility is that SUSD4 binds directly to GLUA2 subunits to promote AMPA receptors degradation and that the SUSD4 interactors modulate this function. In particular, the NEDD4 subfamily of HECT ubiquitin ligases is known to ubiquitinate and target for degradation many key signaling molecules, including GLUA1- and GLUA2-containing AMPA receptors (Schwarz et al., 2010; Widagdo et al., 2017). Ubiquitin ligases of the NEDD4 family bind variants of PY motifs on target substrates and adaptors (Chen et al., 2017). However, GLUA1 and GLUA2 subunits lack any obvious motif of this type. In contrast, two potential PY binding sites are present in the intracellular domain of SUSD4 (**Figure 5C**).

To test whether SUSD4 interacts with GLUA2 and how this interaction might be affected by SUSD4 binding to NEDD4 ubiquitin ligases, co-immunoprecipitation experiments were performed on extracts from heterologous HEK293 cells transfected with SEP-tagged GLUA2 and various HA-tagged SUSD4 constructs (**Figure 5C and 5D**). GLUA2 was detected in extracts obtained after affinity-purification of the HA-tagged full length SUSD4 (HA-SUSD4), while it was absent if HA-SUSD4 was replaced by a control transmembrane protein, PVRL3*α* (**Figure 5D**). To test which domain in SUSD4 is responsible for this interaction, several deletion constructs were generated (**Figure 5C)**. Strong co-immunoprecipitation of GLUA2 was detected in affinity-purified extracts from cells expressing the HA-tagged extracellular domain of SUSD4 alone (HA-SUSD4-N_T_ construct), showing that this domain is sufficient for GLUA2 interaction. Deletion of the extracellular domain (HA-SUSD4*Δ*N_T_) or of the cytoplasmic domain (HA-SUSD4*Δ*C_T_) of SUSD4 did not abrogate binding to GLUA2, showing the cooperation of several domains of SUSD4 for the binding to the GLUA2 subunit (**Figure 5D**). Furthermore, the level of GLUA2 recovered in extracts from HA-SUSD4*Δ*C_T_ transfected cells was intermediate between the one affinity-purified from HA-SUSD4-N_T_ and HA-SUSD4 transfected cells, suggesting that the cytoplasmic domain of SUSD4 plays a regulatory role for GLUA2 binding.

To test whether the two PY motifs in the intracellular tail of SUSD4 mediate the binding of NEDD4 ubiquitin ligases and whether mutations in these sites could change SUSD4’s ability to interact with GLUA2, we generated single- and double-point mutants. While the mutation of the PPxY site (SUSD4-*Δ*PY mutant) abrogated binding of SUSD4 only partially, mutation of only the LPxY site (SUSD4-*Δ*LY mutant) or of both sites (SUSD4-*Δ*PY/LY mutant) completely prevented the binding to NEDD4 ubiquitin ligases (**Figures 5C and D**). These mutations did not change significantly the level of SUSD4 protein in transfected HEK293 cells suggesting that the degradation of SUSD4 itself is not regulated by NEDD4 ubiquitin ligases (**Figure S11**). The same levels of SEP-GLUA2 were detected in affinity-purified samples whether HA-tagged full-length SUSD4 or the point mutants unable to bind NEDD4 ubiquitin ligases were used as a bait (**Figure 5D**). This shows that interaction of NEDD4 ubiquitin ligases does not affect SUSD4’s ability to interact with GLUA2.

Overall, our results suggest that the interaction of SUSD4 with GLUA2 could directly promote its activity-dependent degradation and that this function could be regulated by several molecular partners (**Figure 5E)**. In particular, direct interaction with NEDD4 ubiquitin ligases could either promote GLUA2 ubiquitination in an activity-dependent manner in neurons or regulate the interaction of SUSD4 with other pathways revealed by our proteomics analysis (**Figure 5E**).

## Discussion

Our study shows that the CCP domain-containing protein SUSD4 starts to be expressed in various neurons of the mammalian central nervous system when synapses are formed and mature. *Susd4* loss-of-function in mice leads to impaired motor coordination adaptation and learning, misregulation of synaptic plasticity in cerebellar PCs and deficient activity-dependent degradation of GLUA2-containing AMPA receptors. SUSD4 can bind GLUA2 subunit and controls its degradation. This function of SUSD4 could be regulated by its direct binding to ubiquitin ligases of the NEDD4 family.

### SUSD4 promotes long-term synaptic depression

The choice between recycling of AMPA receptors to the membrane or targeting to the endolysosomal compartment for degradation is key for the regulation of the number of AMPA receptors at synapses, as well as for the direction and degree of activity-dependent synaptic plasticity (Ehlers, 2000; Lee et al., 2002). Blocking trafficking of AMPA receptors through recycling endosomes, for example using a RAB11 mutant, prevents long-term potentiation (LTP) in neurons (Park et al., 2004). Conversely, blocking the sorting of AMPA receptors to the endolysosomal compartment, for example using a RAB7 mutant, impairs long-term depression (LTD) in hippocampal CA1 pyramidal neurons and cerebellar Purkinje cells (PCs) (Fernandez-Monreal et al., 2012; Kim et al., 2017). Further support for the role of receptor degradation comes from mathematical modeling showing that in cerebellar PCs LTD depends on the regulation of the total pool of glutamate receptors (Kim et al., 2017). The GLUA2 AMPA receptor subunit, and its regulation, is of particular importance for LTD (Diering and Huganir, 2018). Phosphorylation in its C-terminal tail and the binding of molecular partners such as PICK1 and GRIP1/2 is known to regulate endocytosis and recycling (Bassani et al., 2012; Chiu et al., 2017; Fiuza et al., 2017), and mutations in some of the phosphorylation sites leads to impaired LTD (Chung et al., 2003). The molecular partners regulating the targeting for degradation of GLUA2 subunits in an activity-dependent manner during LTD remain to be identified. Our study shows that loss-of-function of *Susd4* leads both to loss of LTD and loss of activity-dependent degradation of GLUA2 subunits. Loss-of-function of *Susd4* does not affect degradation of another postsynaptic receptor, GluD2, showing the specificity of SUSD4 action. Furthermore, loss-of-function of *Susd4* facilitates LTP of PF/PC synapses. Overall our results support a model in which, during LTD, a specific molecular machinery containing SUSD4 promotes targeting of GLUA2-containing AMPA receptors to the degradation compartment in an activity-dependent manner, away from the recycling pathway that promotes LTP.

### SUSD4 might promote GLUA2 subunits targeting to the degradation compartment

The degradation of specific targets such as neurotransmitter receptors must be regulated in a stimulus-dependent and synapse-specific manner in neurons, to ensure proper long-term synaptic plasticity, learning and memory (Tai and Schuman, 2008). How is this level of specificity achieved? Adaptor proteins, such as GRASP1, GRIP1, PICK1 and NSF, are known to promote AMPA receptor recycling and LTP (Anggono and Huganir, 2012). Such adaptors for the promotion of LTD remain to be found.

Our results show that SUSD4 and GLUA2 AMPA receptor subunits interact in cells, colocalize in neurons, and that SUSD4 promotes GLUA2 degradation. Furthermore, our affinity-purification experiments identified SUSD4 as a binding partner for HECT E3 ubiquitin ligases of the NEDD4 family. The family of HECT E3 ubiquitin ligases contains 28 enzymes including the NEDD4 subfamily that is characterized by an N-terminal C2 domain, several WW domains and the catalytic HECT domain (Weber et al., 2019). This subgroup of E3 ligases adds K63 ubiquitin chains to their substrate, a modification that promotes sorting to the endolysosomal compartment for degradation (Boase and Kumar, 2015). NEDD4 E3 ligases are highly expressed in neurons in the mammalian brain and have many known substrates with various functions, including ion channels and the GLUA1 AMPA receptor subunit. Accordingly, knockout mice for the *Nedd4-1* gene die during late gestation (Kawabe et al., 2010). The activity and substrate selectivity of NEDD4 E3 ligases thus need to be finely tuned. Both GLUA1 and GLUA2 AMPA receptor subunits are ubiquitinated on lysine residues in their intracellular tails in an activity-dependent manner (Lin et al., 2011; Lussier et al., 2011; Schwarz et al., 2010; Widagdo et al., 2015). Mutation of these lysine residues decreases localization of GLUA1 and GLUA2 AMPA receptor subunits in the endolysosomal compartment in neurons (Widagdo et al., 2015). However, GLUA1 and GLUA2 subunits lack any obvious intracellular direct binding motif to the WW domain of NEDD4 ubiquitin ligases, raising questions about the precise mechanism allowing regulation of AMPA subunits trafficking and degradation by these enzymes. SUSD4 could play a role in regulating targeting of NEDD4 ubiquitin ligases to AMPA receptors in an activity-dependent manner in neurons. Alternatively, the interaction of SUSD4 with NEDD4 ubiquitin ligases might regulate the trafficking of the SUSD4/GLUA2 complex to the degradation pathway.

Further work is warranted to determine the mechanism of action of SUSD4 in neurons. In particular how is the specificity of its action achieved and how is neuronal activity regulating this pathway? Among the potential partners of SUSD4 identified by our proteomics analysis, several encompass functions that are relevant for the regulation of synaptic plasticity, such as receptor anchoring, clathrin mediated endocytosis and proteasome function **(Figure 5E)**. Moreover, our results suggest that not all spines contain SUSD4 and thus an attractive hypothesis is that its recruitment to synapses is activity-dependent. However another possibility lies in activity-dependent proteolysis, which has been shown to regulate the function of several types of cell adhesion molecules (Shinoe and Goda, 2015). The presence of four CCP domains in the extracellular region of SUSD4 suggests that it could bind proteins extracellularly. In particular, it was previously shown that SUSD4 can bind the C1Q globular domain of the complement protein C1 (Holmquist et al., 2013), a domain that is also found in presynaptic proteins of the C1Q family known for their role in synapse formation and function (Sigoillot et al., 2015; Südhof, 2018; Yuzaki, 2011). SUSD4 could thus enable fine spatiotemporal regulation of the degradation of GLUA2-containing AMPA receptors in a trans-synaptic manner.

### SUSD4 and neurodevelopmental disorders

In humans, the 1q41-42 deletion syndrome is characterized by many symptoms including IDs and seizures, and in a high majority of the cases the microdeletion encompasses the *Susd4* gene (Rosenfeld et al., 2011). A *Susd4* copy number variation has been identified in a patient with autism spectrum disorder (ASD)(Cuscó et al., 2009). *Susd4* was recently identified amongst the 124 genes with genome wide significance for *de novo* mutations in a cohort of more than 10,000 patients with ASD or IDs (Coe et al., 2019). The *GRIA2* gene (coding for the GLUA2 subunit) has been found as an ASD susceptibility gene (Salpietro et al., 2019; Satterstrom et al., 2018) and mutations or misregulation of ubiquitin ligases have been found in many models of ASDs or intellectual deficiencies (Cheon et al., 2018; Lee et al., 2018; Satterstrom et al., 2018). For example, ubiquitination of GLUA1 by NEDD4-2 is impaired in neurons from a model of Fragile X syndrome (Lee et al., 2018). Understanding the precise molecular mechanisms underlying the activity-dependent degradation of GLUA2 by the SUSD4/NEDD4 complex will thus be of particular importance for our understanding of the etiology of these neurodevelopmental disorders.

Mutations in the *Susd4* gene might contribute to the etiology of neurodevelopmental disorders by impairing synaptic plasticity. Deficits in LTD such as the one found in the *Susd4* KO mice are a common feature of several mouse models of ASDs (Auerbach et al., 2011; Baudouin et al., 2012; Piochon et al., 2014). *Susd4* loss-of-function leads to motor impairments, a symptom that is also found in ASD patients (Fournier et al., 2010). Very recently, a reduction in exploratory behavior, in addition to impairments of motor coordination, was reported after *Susd4* loss-of-function (Zhu et al., 2020). Long-term synaptic plasticity has been proposed as a mechanism for learning and memory. While *in vivo* evidence for the role of LTP in these processes has accumulated, the role of LTD is still discussed (Andersen et al., 2017; Kakegawa et al., 2018; Raymond and Medina, 2018; Schonewille et al., 2011). Because of the broad expression of SUSD4 and of ubiquitin ligases of the NEDD4 subfamily in the mammalian central nervous system, this pathway is likely to control synaptic plasticity at many synapse types and its misregulation might lead to impairments in many learning and memory paradigms.

## Acknowledgments

We gratefully acknowledge the Collège de France imaging facility (IMACHEM-IBiSA), in particular P. Mailly for help with the design of the macro for GLUA2 quantification and Estelle Anceaume for help with image acquisition. We also thank the personnel from the CIRB, INCI and chronobiotron CNRS UMS 3415, IBPS and IBENS animal facilities. We would like to thank Philippe Marin for advice on proteomics analysis. Mass spectrometry experiments were carried out using facilities of the Functional Proteomics Platform of Montpellier.

## Contributions

FS, PI, LR-R, IG-C and KI designed the study and the experiments. IG-C, KI, MC, AK, FAG, JV, MS, ST, MA, OV, YN, AD, FS, SMS, MV and CB-L performed the experiments, and collected the data; AT and J-LB provided the SUSD4 knockout mice and conceptual advice; FS and IG-C wrote the first draft of the manuscript; all authors read the manuscript and KI, MC, AK, MS, SMS, CB-L, MV, AD, J-LB, PI and LR-R revised the manuscript.

## Data and materials availability

All data is available in the main text or the supplementary materials.

## Supplementary Information

Materials and Methods

Figs. S1 to S11

Tables S1

## Supplementary Information

▪ Table S1
▪ Figures S1 to S11
▪ Materials and Methods
▪ Supplementary references

**Supplementary Table 1.** Behavioral characterization of *Susd4* KO mice. From three month-old *Susd4* knockout (KO) and *wild type* (WT) littermates. Mean ± s.e.m. or percentage of mice (Physical Characteristics: WT n=10 and KO n=12 mice; Sensorimotor Reflexes and Motor responses: WT n=24 and KO n=24 mice).

|  | WT |  | Susd4 KO |  |
| --- | --- | --- | --- | --- |
| *Physical Characteristics* |  |  |  |  |
| Weight (g) | 24,13 | ± 1.20 | 24,72 | ± 1.33 |
| Whiskers (% with) | 80 | % | 83,3 | % |
| Palpebral Closure (% with) | 0 | % | 0 | % |
| Piloerection (% with) | 20 | % | 25 | % |
| *Sensorimotor Reflexes* |  |  |  |  |
| *(% subjects displaying "normal response")* |  |  |  |  |
| Cage movement | 100 | % | 100 | % |
| Whisker response | 100 | % | 100 | % |
| Eye Blink | 100 | % | 100 | % |
| Ear Twitch | 100 | % | 100 | % |
| *Motor Responses* |  |  |  |  |
| Open Field Locomotion |  |  |  |  |
| Improvement (number) | 22.83 | ± 2,89 | 19,42 | ± 2,03 |
| Distance (cm) | 2764 | ± 235.0 | 2301 | ± 158,5 |
| Speed (cm/s) | 13.32 | ± 0,42 | 13,06 | ± 0,30 |
| Time on Center (%) | 13.30 | ± 1,33 | 11,18 | ± 1,26 |

**Figure Supplementary 1.**
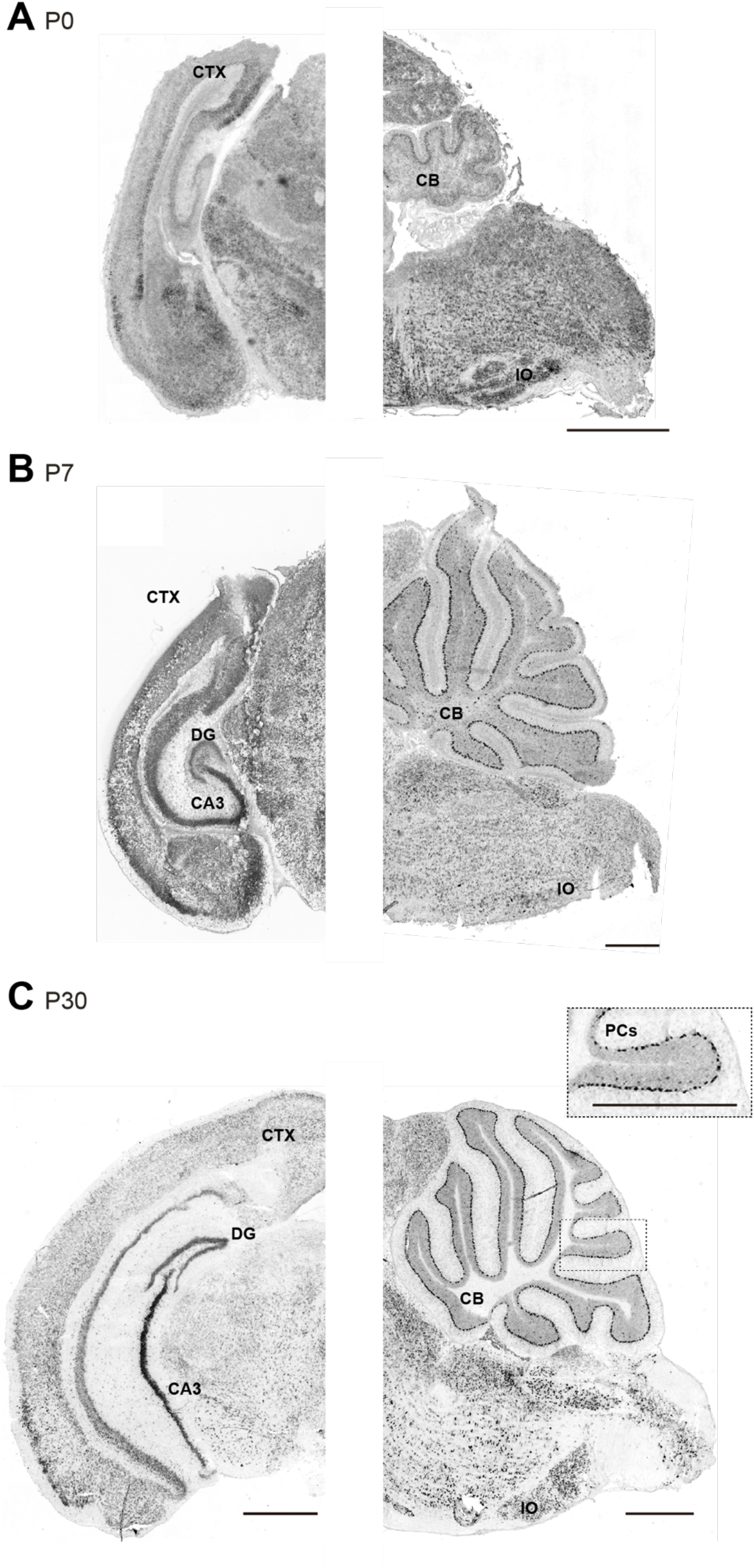
*Susd4* mRNA expression in the developing mouse brain. **(A)** *Susd4* mRNA expression was visualized in the brain of wild-type (WT) mice by *in situ* hybridization. Coronal (left) and sagittal (right) sections are presented at postnatal day 0 (P0), **(B)** postnatal day 7 (P7) and **(C)** postnatal day 30 (P30). *Susd4* expression was found in many regions including the cerebral cortex (CTX), the dentate gyrus (DG) and CA3 regions in the hippocampus (coronal section, left), the cerebellum (CB), in particular Purkinje cells (PCs), and the inferior olive (IO; sagittal section, right). Scale bars: 250µm and 500µm (inset **C**).

**Figure Supplementary 2.**
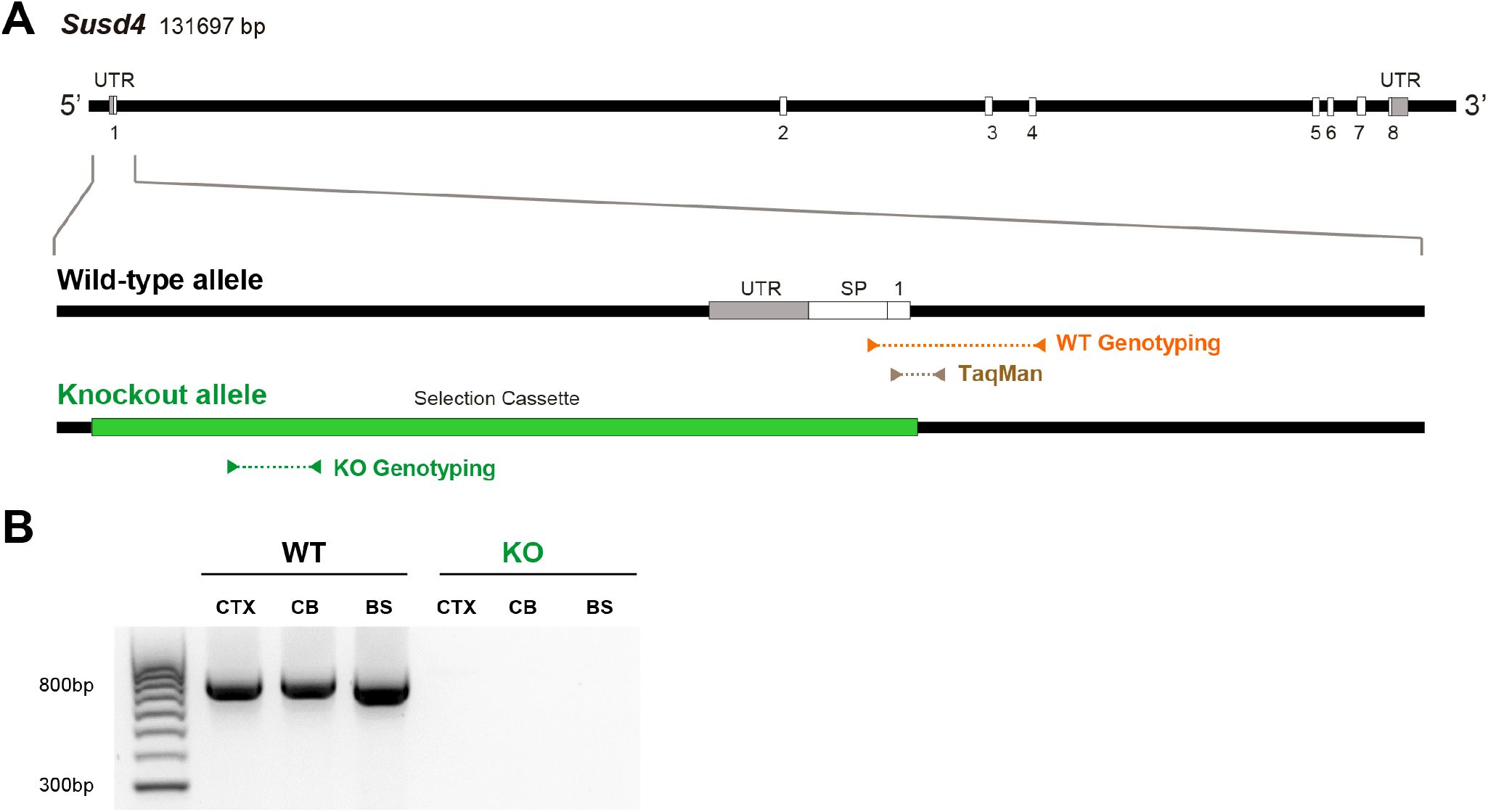
Characterization of *Susd4* knockout (KO) mice. **(A)** Structure of the *Susd4* gene and strategy for the generation of the knockout mouse. The gene coding for the *Susd4* mRNA contains 8 exons. The wild-type WT allele is presented indicating the localization of the primers used for genotyping and of the probes used for TaqMan RT-qPCR. In the knockout allele, the 5’UTR and first exon are entirely deleted and replaced by the selection cassette. **(B)** *Susd4* expression was assessed by RT-PCR using primers encompassing exons 6 to 8 in extracts from cortex (CTX), cerebellum (CB) and brainstem (BS) in control and *Susd4* KO mice.

**Figure Supplementary 3.**
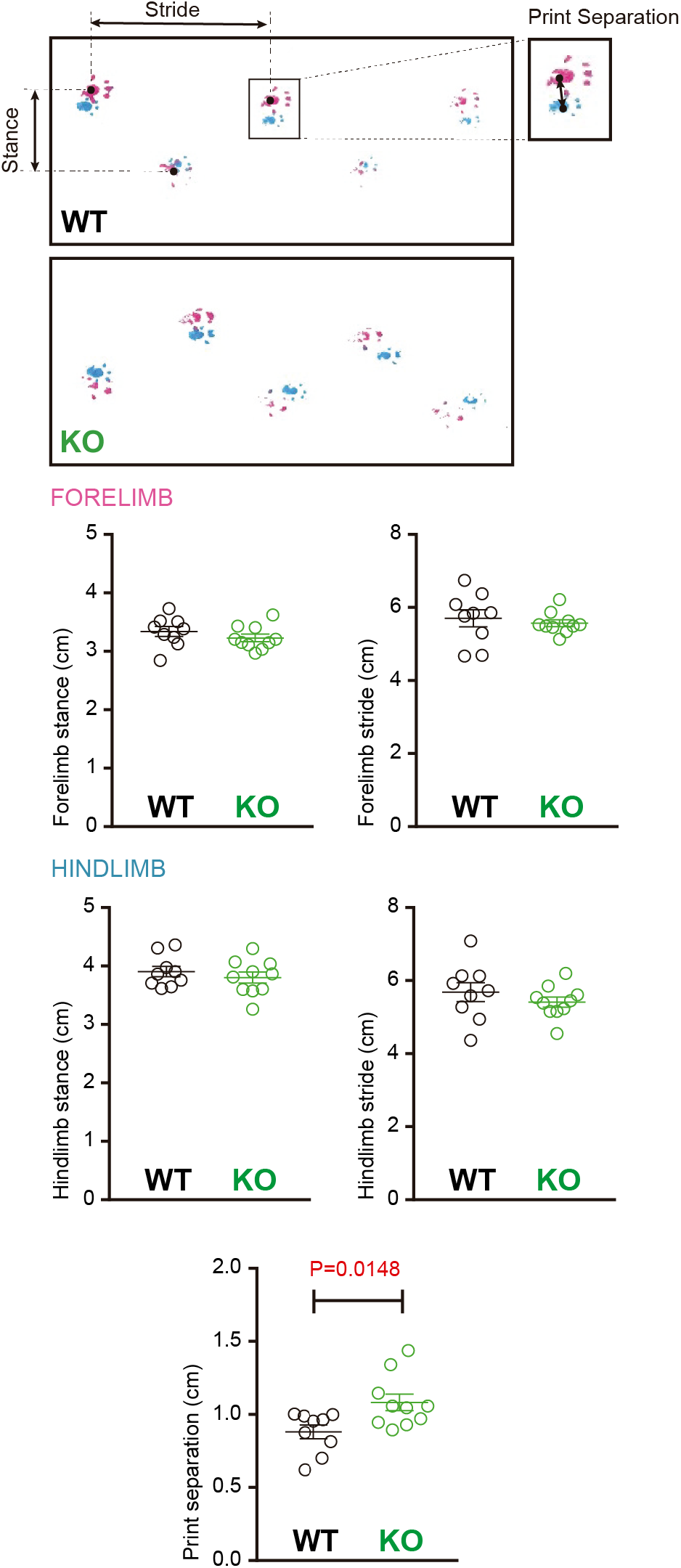
Footprint analysis in *Susd4* KO mice. Footprint patterns of P30 WT and *Susd4* KO mice were quantitatively analyzed by measuring stride length for the fore paws (magenta) and hind paws (cyan), stance length for the forelimbs and hindlimbs, and print separation. Mean ± s.e.m. (WT n=9 and KO n=10 mice; unpaired Student’s t-test; Forelimb stance: P=0.3059; Forelimb stride: P=0.5882; Hindlimb stance: P=0.4533; Hindlimb stride: P=0.3580; Print separation: * P=0.0148).

**Figure Supplementary 4.**
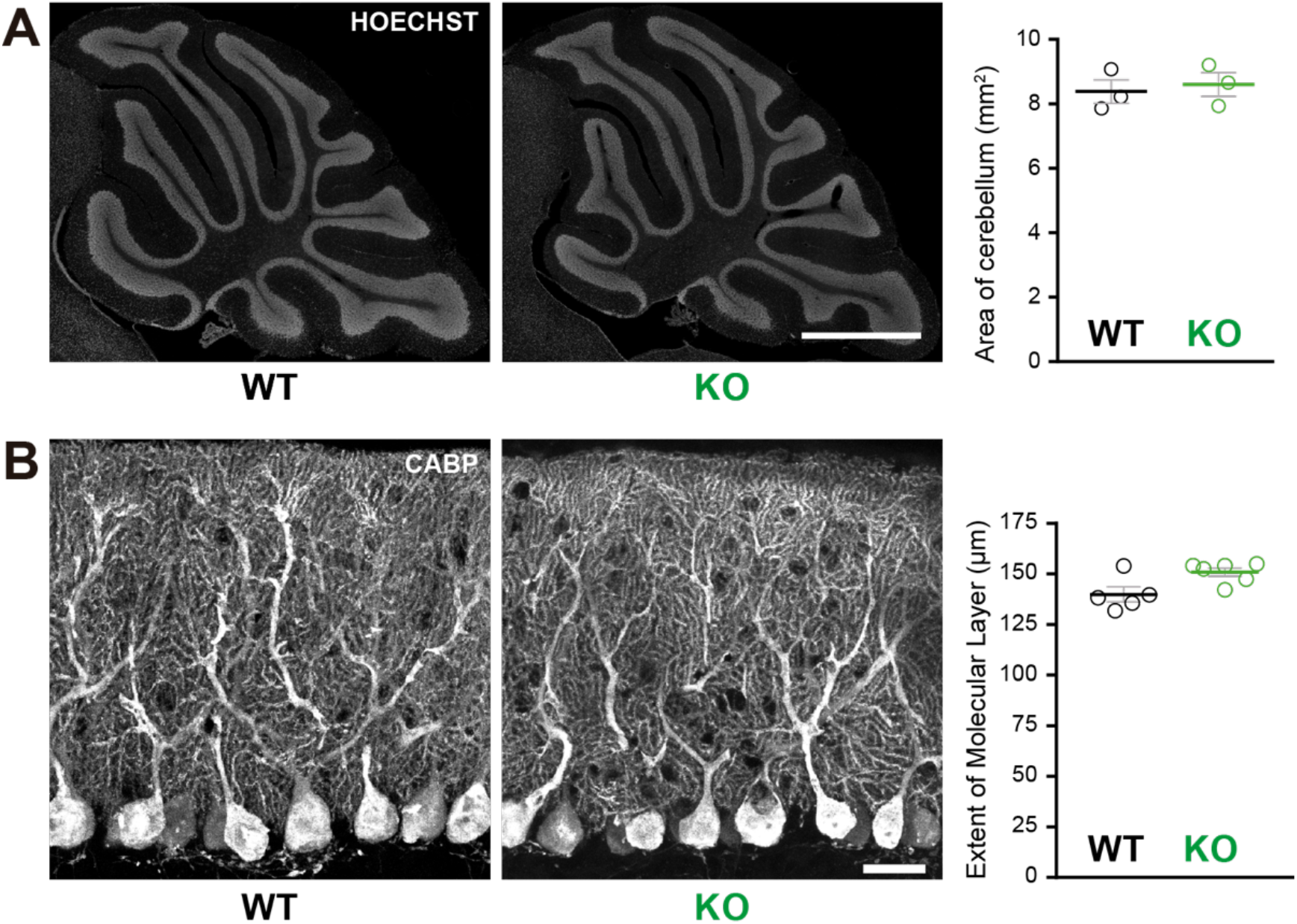
Normal cerebellar cytoarchitecture in *Susd4* KO mice. **(A)** Parasagittal sections of P30 WT and *Susd4* KO cerebella were stained with Hoechst and used for quantitative analysis of the mean area of the cerebellum. Mean ± s.e.m. (n=3 WT mice and n=3 KO mice). Scale bar: 500µm. **(B)** Calbindin protein (CABP) immunostaining was used for quantitative analysis of the mean height of the molecular layer. Mean ± s.e.m. (WT n=5 and KO n=6 mice). Scale bar: 30µm.

**Figure Supplementary 5.**
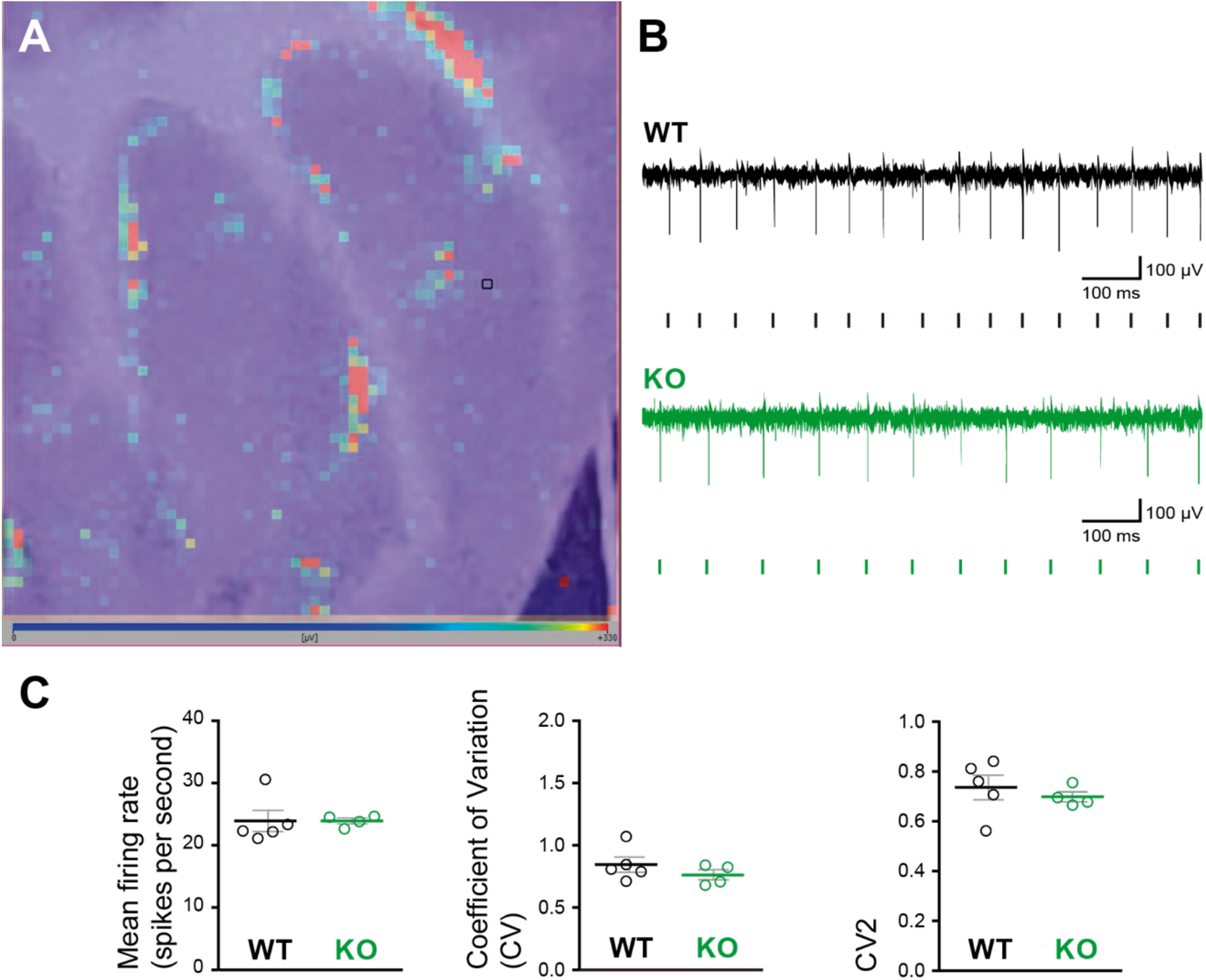
High density microelectrode array (MEA) analysis of Purkinje cell spiking in acute cerebellar slices from *Susd4* KO compared to WT. **(A)** Image of a cerebellar acute slice from a WT mouse overlapped with the image of the color map of the MEA recording. Each pixel represents one channel, where the active units are in red. The black square highlights one of the channels. **(B)** Representative traces of electrical activity recorded in one channel from control and *Susd4* KO mice. Each tick points out one action potential that has been detected and sorted by the Brainwave software. **(C)** Histograms of the mean firing rate, coefficient of variation (CV) of Inter Spike Intervals and CV2. Mean ± s.e.m. (WT n=5 and KO n=4 mice; Mann Whitney test; Mean Firing Rate: P=0.2857; CV: P=0.4127; CV2: P=0.5373).

**Figure Supplementary 6.**
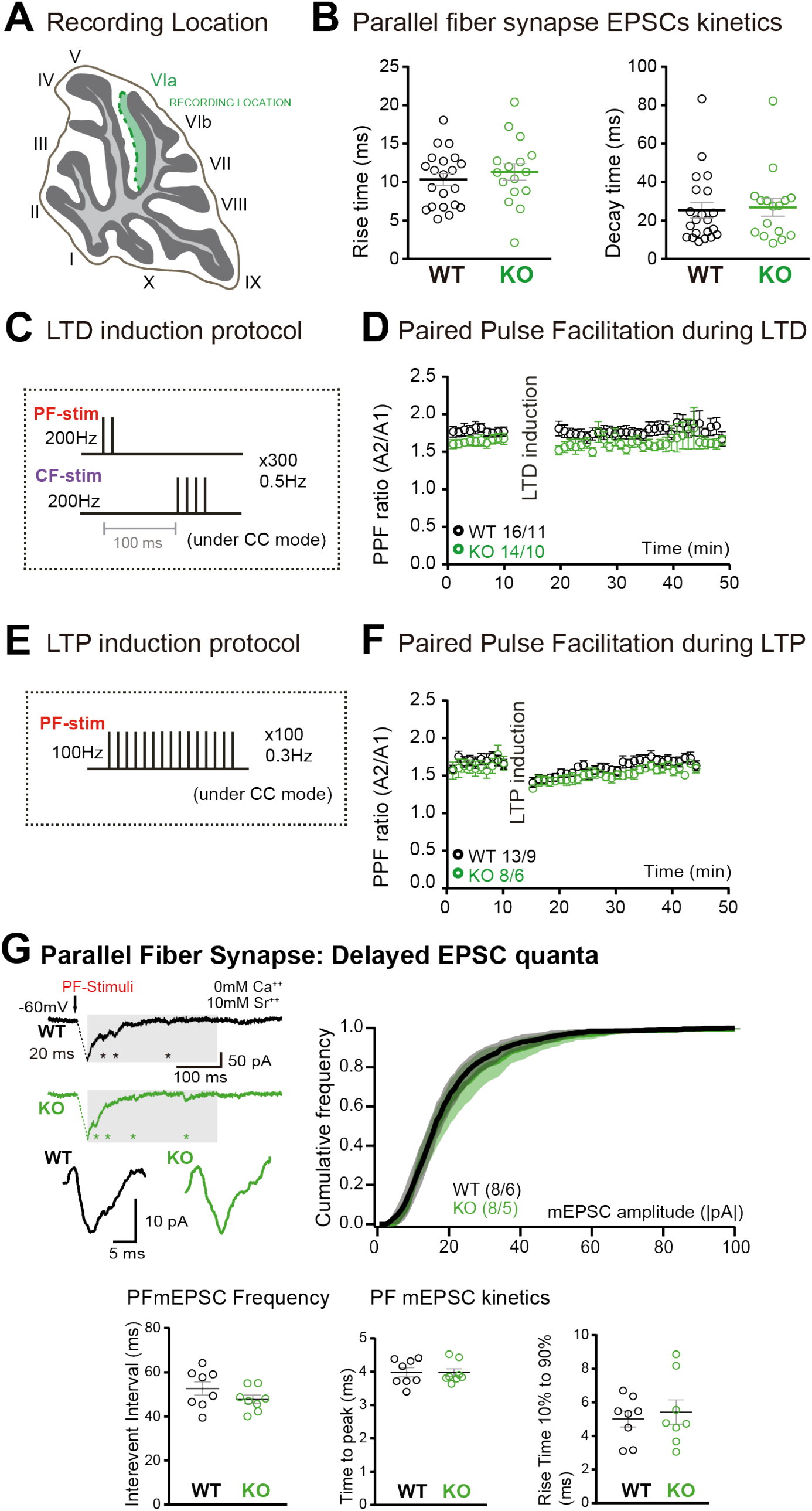
Parallel fiber (PF) /Purkinje cell (PC) synapse EPSCs kinetics, long-term plasticity induction protocols, paired-pulse facilitation ratio and delayed EPSC quanta. **(A)** Schematic representation of the recording location (internal lobule VIa of the vermis). **(B)** No change in the rise time and decay of Parallel fiber/Purkinje cell EPSCs was induced by *Susd4* deletion. Mean ± s.e.m. (WT n=21 cells from 8 mice and KO n=16 cells from 6 mice; Rise time: unpaired Student’s t-test, P=0.4570; Decay time: Mann Whitney test, P=0.7276). **(C)** Parallel fiber long-term depression (LTD) induction protocol. **(D)** Paired-pulse ratio (A2/A1) during LTD measured at 20 Hz. Mean ± s.e.m. (WT n=16 cells from 11 mice and KO n=14 cells from 10 mice; two-way ANOVA with repeated measures, Interaction (time and genotype): P=0.9935, F(39, 1092)=0.5222). **(E)** Parallel fiber long-term potentiation (LTP) induction protocol. **(F)** Paired-pulse ratio (A2/A1) during LTP measured at 20Hz. Mean ± s.e.m. (WT n=13 cells from 9 mice and KO n=8 cells from 6 mice, two-way ANOVA with repeated measures, Interaction (time and genotype): P=0.9366, F(39, 741)=0.6745). **(G)** Delayed PF-EPSC quanta were evoked by PF stimulation in the presence of strontium (Sr^++^) instead of calcium (Ca^++^) to induce desynchronization of fusion events. Representative sample traces are presented. The cumulative probability for the amplitude shows no difference with *Susd4* loss-of-function. The individual frequency values for each cell (measured as interevent interval) present no differences between the genotypes. No change in the time to peak and in the rise time of PF/PC synapse delayed EPSC quanta was induced by *Susd4* deletion. Mean ± s.e.m. (WT n=8 cells from 6 mice and KO n=8 cells from 5 mice; Amplitude: Kolmogorov-Smirnov distribution test, P=0.1667; Frequency: Mann Whitney test, P=0.1913; Time to peak: Mann Whitney test, P=0.6454; Rise time 10% to 90%: unpaired Student’s t-test, P=0.6486).

**Figure Supplementary 7.**
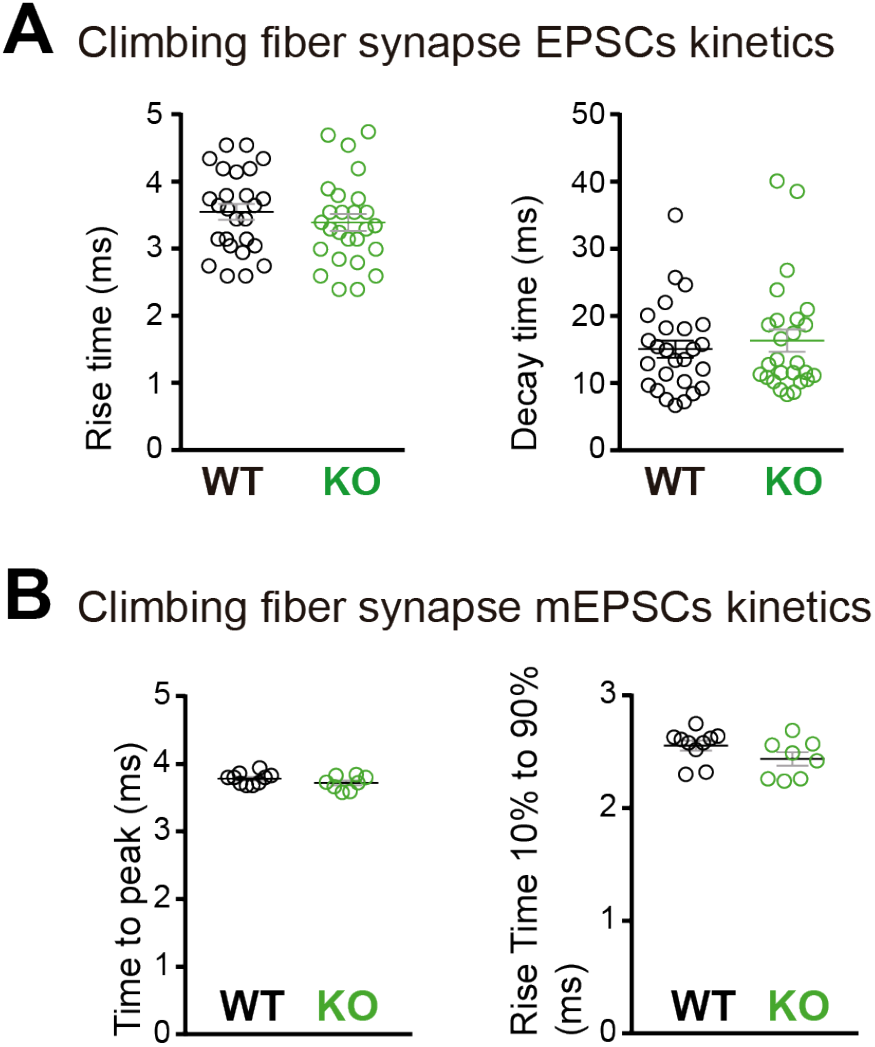
Kinetics of the climbing fiber/Purkinje cell synapse EPSC. **(A)** No change in the rise and decay times of climbing fiber/Purkinje cell EPSCs was induced by *Susd4* deletion. Mean ± s.e.m. (WT n=26 cells from 9 mice and KO n=26 cells from 7 mice; Rise time: unpaired Student’s t-test, P=0.3750; Decay time: Mann Whitney test, P=0.7133). **(B)** No change in the time to peak and in the rise time of CF/PC synapse delayed EPSC quanta was induced by *Susd4* loss-of-function. Mean ± s.e.m. (WT n=10 cells from 6 mice and KO n=8 cells from 3 mice; Time to peak: unpaired Student’s t-test, P=0.1692; Rise time 10% to 90%: Mann Whitney test, P=0.0639).

**Figure Supplementary 8.**
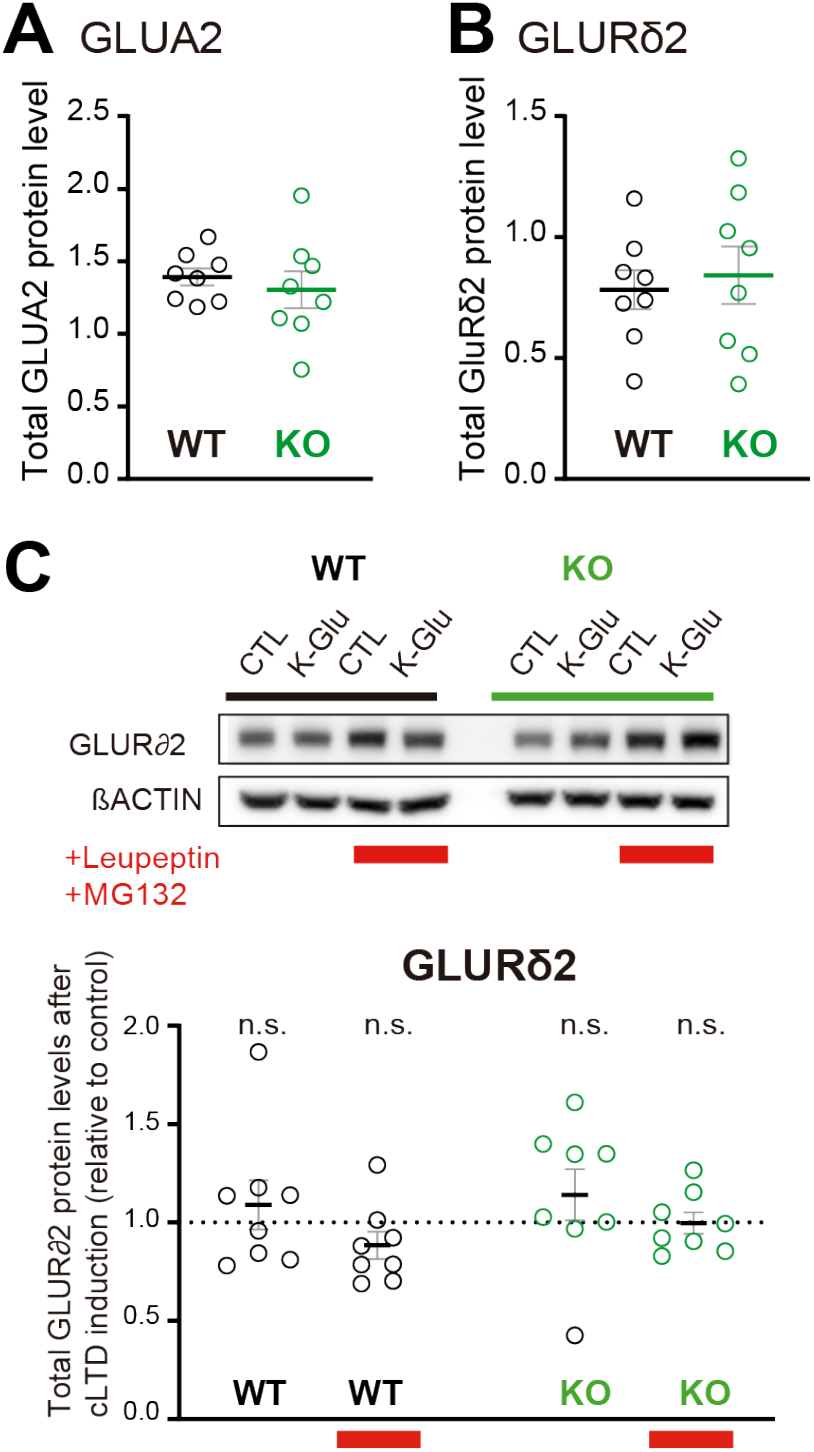
Total GLUA2 and GLURδ2 levels after modulation of SUSD4 expression. **(A)** and **(B)** Total protein levels (normalized to βACTIN) of GLUA2 **(A)** and GLURδ2 **(B)** were not changed in acute cerebellar slices from WT or *Susd4* KO mice in basal conditions. Mean ± s.e.m. (n=8 independent experiments; unpaired Student’s t-test; GLUA2: P=0.5424; GLURδ2: P=0.6821). **(C)** Cerebellar acute slices from control WT and *Susd4* KO mice were incubated to induce chemical LTD (cLTD; K-Glu: K^+^ 50mM and glutamate 10µM for 5min followed by 30min of recovery). Slices were incubated with 100µg/mL leupeptin and with 50µM MG132 (to inhibit lysosomal and proteasome degradation, respectively). Band intensities of GLURδ2 were normalized to βACTIN. The ratios between levels with cLTD induction (K-Glu) and without cLTD induction (CTL) are represented. Mean ± s.e.m. (n=8 independent experiments; two-tailed Student’s one sample t-test was performed on the ratios with a null hypothesis of 1, P_WT_ = 0.4973, P_WT+Leu/MG132_ = 0.1433, P_KO_ = 0.3143, P_KO+Leu/MG132_ = 0.9538, n.s.= not significant).

**Figure Supplementary 9.**
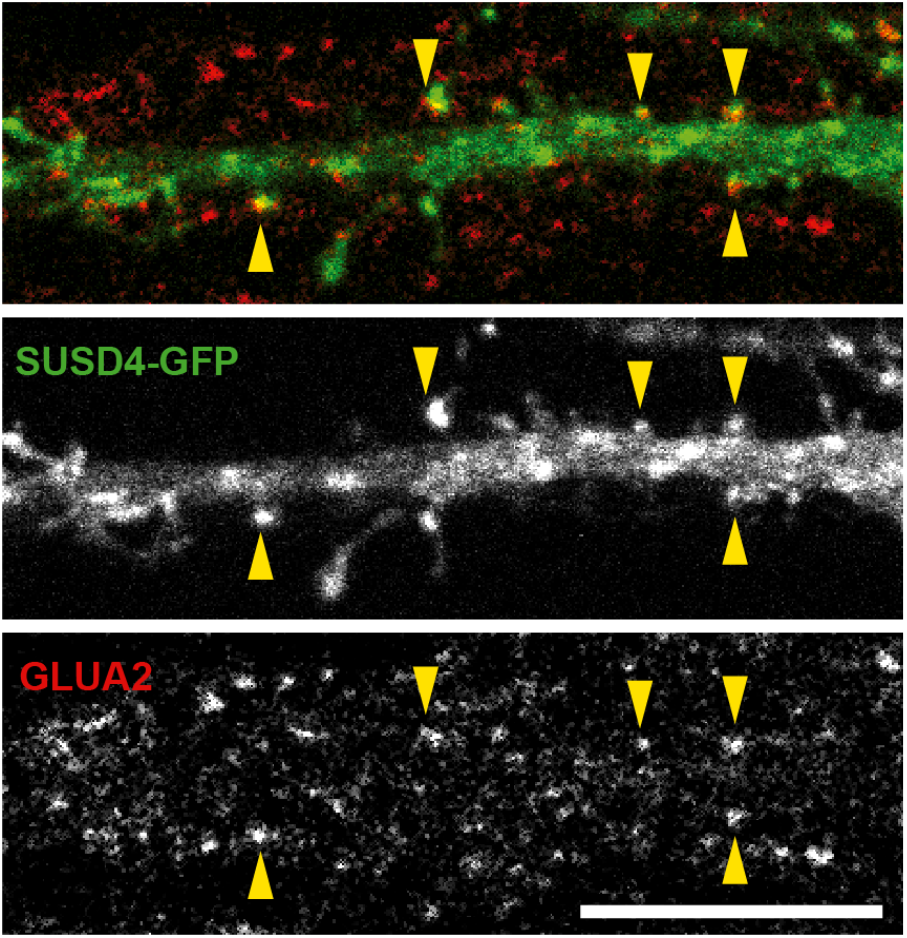
HA-SUSD4 colocalizes with AMPA receptor subunit GLUA2 in hippocampal neurons. Mouse hippocampal neurons were transfected at 13 days *in vitro* (DIV13) with a GFP-tagged SUSD4 construct and immunostained at DIV17 for green fluorescent protein (GFP, green) to localize SUSD4 and for the endogenous GLUA2 subunit (anti-GLUA2, red). The arrowheads indicate the spines containing SUSD4 and GLUA2. Scale bar: 10 µm.

**Figure Supplementary 10.**
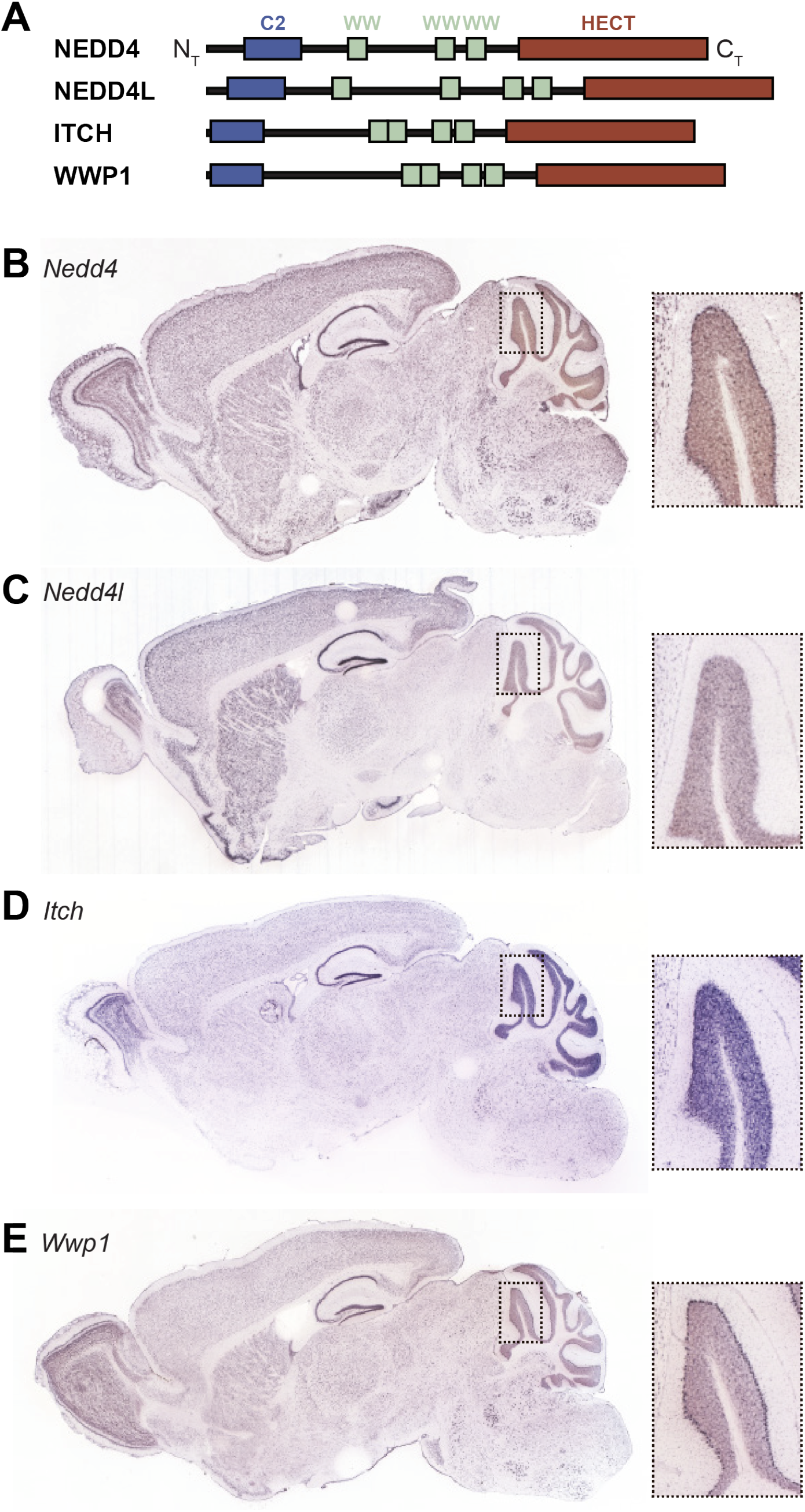
Expression of HECT ubiquitin ligases in adult mouse brain. **(A)** Schematic representation of four SUSD4 interactors: NEDD4, NEDD4L, ITCH and WWP1. Legends: N_T_, N-terminus; HECT, Homologous to the E6-AP C-terminus domain; C_T_, C-terminus. **(B)** Pattern of expression of *Nedd4* (RP_050712_03_C08), **(C)** *Nedd4l* (RP_040625_01_G10), **(D)** *Itch* (RP_050222_01_H06) and **(E)** *Wwp1* (RP_050510_02_E12) mRNA in the adult mouse brain. From Allen Brain Atlas (www.brain-map.org).

**Figure Supplementary 11.**
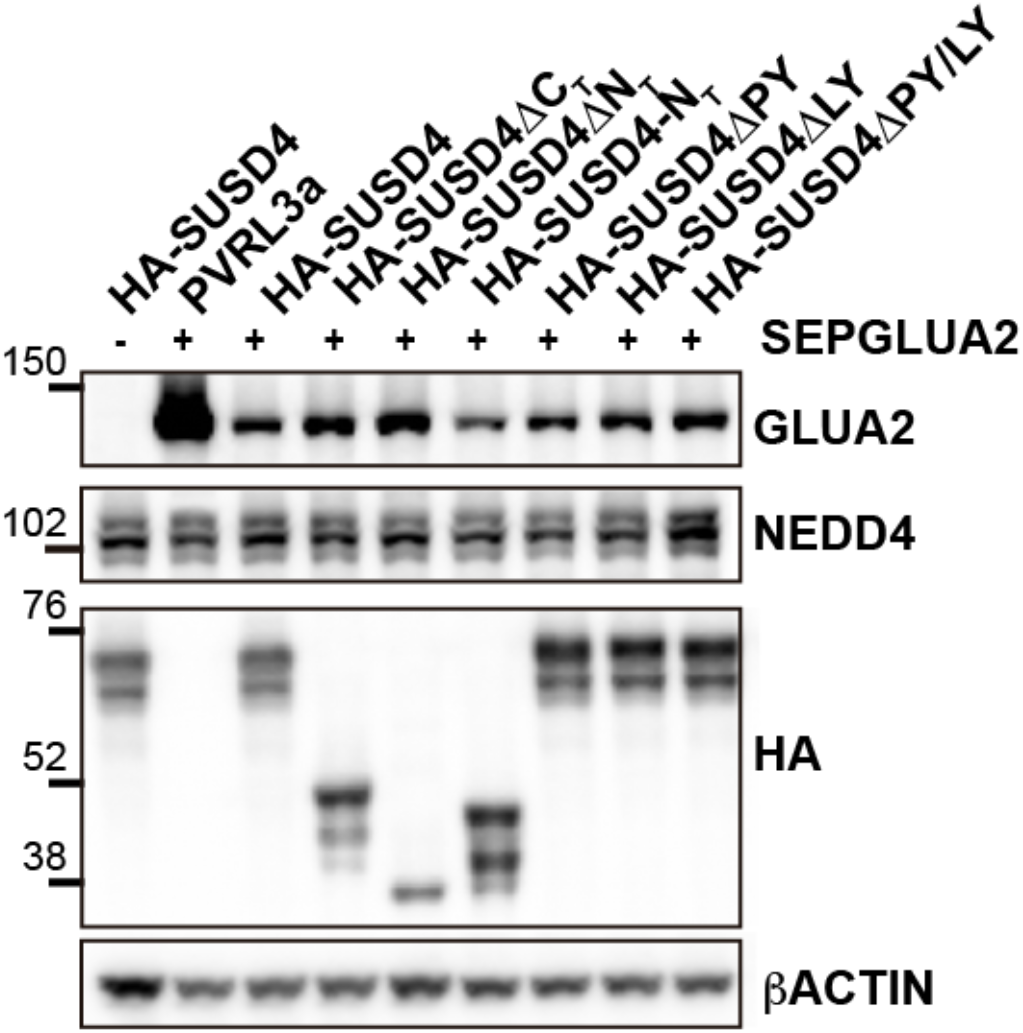
Total protein levels in HEK293 cells transfected with SEP-GLUA2 and different SUSD4 mutant constructs (related to figures 5C and 5D) HEK293 cells were transfected with SEP-GLUA2 together with PVRL3α as a control or one of the HA-SUSD4 constructs for coimmunoprecipitation experiments. Input extracts were probed for GLUA2 (with an anti-GLUA2 antibody), the HECT ubiquitin ligase NEDD4 (anti-NEDD4 antibody), and the HA-tagged SUSD4 constructs (anti-HA antibody). b-ACTIN was used as a loading control. Representative image of N=3 independent experiments.

## Materials and Methods

### Animals

*<u>Susd4</u>* <u>knockout (KO) mice</u> were generated and maintained on the C57BL/6J background (generated by Lexicon Genetics Incorporated, The Woodlands, USA)(Tang et al., 2010). Out of the 8 *Susd4* exons, coding exon 1 (NCBI accession NM_144796.2) and the 5’UTR (NCBI accession BM944003) were targeted by homologous recombination. This resulted in the deletion of a 1.3kb sequence spanning the transcription initiation site and exon 1 (**Figure 1E** and **Supplementary Figure 2A**). Subsequent genotyping of mice was performed using PCR to detect the wild-type (WT) allele (forward primer: 5’ CTG TGG TTT CAA CTG GCG CTG TG 3’; reverse primer: 5’ GCT GCC GGT GGG TGT GCG AAC CTA 3’) or the targeted allele (forward primer: 5’ TTG GCG GTT TCG CTA AAT AC 3’; reverse primer: 5’ GGA GCT CGT TAT CGC TAT GAC 3’). Heterozygous *Susd4*^+/-^ mice were bred to obtain all the genotypes needed for the experiments (*Susd4*^+/+^ (WT) and *Susd4*^-/-^ (KO) mice) as littermates.

The <u>Htr5b-GFP mouse line</u> was used for labeling of climbing fibers (CF; The Gene Expression Nervous System Atlas (GENSAT) Project, NINDS Contracts N01NS02331 & HHSN271200723701C to The Rockefeller University (New York, NY)). Genotyping was performed using the following primers: 5’ TTG GCG CGC CTC CAA CAG GAT GTT AAC AAC 3’ and 5’ CGC CCT CGC CGG ACA CGC TGA AC 3’ (**Figure Supplementary 2A**).

The <u>L7Cre mouse line</u> was obtained from Jackson laboratories (B6.129-Tg(Pcp2-cre)2Mpin/J ; Stock Number: 004146) and genotyping was performed using the following primers: 5’ GGT GAC GGT CAG TAA ATT GGA C 3’; 5’ CAC TTC TGA CTT GCA CTT TCC TTG G 3’ and 5’ TTC TTC AAG CTG CCC AGC AGA GAG C 3’.

All animal protocols were approved by the *Comité Regional d’Ethique en Experimentation Animale* (no. 00057.01) and animals were housed in authorized facilities of the CIRB (# C75 05 12).

### Antibodies

The following primary antibodies were used: mouse monoclonal anti-CABP (1:1000; Swant, Switzerland, Cat#300), rabbit polyclonal anti-CABP (1:1000; Swant, Cat#CB38), mouse monoclonal anti-GFP (1:1000; Abcam, Cambridge, United Kingdom, Cat#ab1218), rabbit polyclonal anti-GFP (1:1000; Abcam, Cat#ab6556), mouse monoclonal anti-GLUA2 (clone 6C4; 1:500; Millipore, Massachusetts, USA, Cat#MAB397 and BD, New Jersey, USA, Cat#556341), rabbit monoclonal anti-GLUA2 (1:1000; Abcam, Cat#ab206293), rabbit polyclonal anti-GLURδ1/2 (1:1000; Millipore, Cat#AB2285), rat monoclonal anti-HA (1:1000; Roche Life Science, Penzberg, Germany, Cat#11867423001), rabbit monoclonal anti-ITCH (1:1000; Cell Signaling Technology, Massachusetts, USA, Cat#12117), rabbit polyclonal anti-NEDD4 (1:10000; Millipore, Cat#07-049), mouse anti-ubiquitin proteins antibody (clone FK2; Enzo, New York, USA, Cat#BML-PW8810 and Sigma, Gothenburg, Sweden, Cat#04-263), guinea pig polyclonal anti-VGLUT1 (1:5000; Millipore, Cat#AB5905), guinea pig polyclonal anti-VGLUT2 (1:5000; Millipore, Cat#AB2251) and rabbit polyclonal anti-WWP1 (1:2000; Proteintech, Chicago, USA, Cat#13587-1-AP).

The following secondary antibodies were used: donkey polyclonal anti-Goat Alexa Fluor 568 (1:1000; Invitrogen, California, USA, Cat#A11057), donkey anti-Mouse Alexa Fluor 488 (1:1000; Invitrogen, Cat#R37114), donkey polyclonal anti-Mouse Alexa Fluor 568 (1:1000; Invitrogen, #A10037), donkey polyclonal anti-Rabbit Alexa Fluor 488 (1:1000; Invitrogen, Cat#A21206), donkey polyclonal anti-Rat Alexa Fluor 594 (1:1000; Invitrogen, #A21209), donkey polyclonal anti-Rat Alexa Fluor 568 (1:1000; Abcam, Cat#175475), goat polyclonal anti-Guinea Pig Alexa Fluor 488 (1:1000; Invitrogen, Cat#A110-73), goat polyclonal anti-Guinea Pig Alexa Fluor 647 (1:1000; Invitrogen, Cat#A21450), goat polyclonal anti-Mouse HRP (1:10000; Jackson Immune Research Laboratories, Pennsylvania, USA, Cat#115-035-174), goat polyclonal anti-rat HRP (1:10000; Jackson Immune Research Laboratories, #112-035-175) and mouse polyclonal anti-rabbit HRP (1:10000; Jackson Immune Research Laboratories, #211-032-171).

The following conjugated antibodies were used: sheep polyclonal anti-digoxigenin alkaline phosphatase (1:2000 - 1:5000; Roche Life Science, Cat#11093274910), mouse monoclonal anti-βACTIN (clone AC-15) HRP (1:25000; Abcam, Cat#ab49900), rabbit polyclonal anti-GFP Alexa Fluor 647 (1:1000; Invitrogen, Cat#A31852), mouse monoclonal anti-GluR2 (clone 6C4) Alexa Fluor 488 (1:1000; Millipore, Cat#MAB397A4) and mouse monoclonal anti-HA (clone 2-2.2.14) DyLight 650 (1:1000; Thermo Fisher Scientific, Massachusetts, USA, Cat#26183-D650).

### Plasmids

Full-length *Susd4* mouse gene was cloned into the mammalian expression vector pEGFP-N1 (Addgene, Massachusetts, USA, Cat#6085-1) to express a SUSD4-GFP fusion construct under the control of the CMV promoter (pSUSD4-GFP). An N-terminal HA tag was inserted just after the signal peptide (pHA-SUSD4-GFP). pHA-SUSD4 was obtained by removal of the C-terminal GFP of pHA-SUSD4-GFP. A truncated form of *Susd4*, expressing the HA-SUSD4-ΔC_T_ mutant, was obtained by inserting a stop codon downstream of the sixth exon, 39bp after the transmembrane domain using PCR on the pHA-SUSD4-GFP plasmid and the following primers: forward primer 5’ GCG CTA GCG ATG TAT CCT TAT GAT GTT CCT G 3’; reverse primer 5’TAG CGG CCG CTA TTA GGG GGG GAA GTG GGC CTT 3’. Other mutant constructs were similarly obtained: the truncated form HA-SUSD4-ΔN_T_ corresponding to aminoacids 294-490, and the extracellular form of SUSD4, HA-SUSD4-N_T_, corresponding to aminoacids 2-299. The HA-SUSD4-ΔPY contains a mutation in aminoacids 411 and 414 changing PPAY to APAA while HA-SUSD4-ΔLY is mutated in aminoacids 376 and 379 changing LPTY to APTA. Mutagenesis was performed using the QuikChange Lightning Multi site directed mutagenesis kit (Agilent, Santa Clara, USA, Cat#210513) according to the manufacturer’s instructions. The plasmid pIRES2-eGFP (Addgene, Cat#6029-1) was used as transfection control. The plasmid expressing SEPGLUA2 (Addgene, Cat#24001) was used to follow GLUA2. The control transmembrane protein PVRL3α was cloned into the mammalian expression vector pCAG-mGFP (Addgene, Cat#14757) to express the protein under the pCAG promoter (pCAG-PVRL3α). The plasmids expressing GFP-tagged RAB proteins (GFP-RAB4a, GFP-RAB5a, GFPRab7a and GFP-RAB11a) were kindly provided by Dr. Bruno Goud.

### Viral mediated *in vivo* expression of HA-SUSD4

AAV2 particles were generated using a hSYN-DIO-HA-SUSD4-2A-eGFP-WPRE construct (Vector biolabs, Malvern, USA) and injected stereotaxically in cerebella of adult mice expressing the CRE recombinase in cerebellar Purkinje cells (PCs) using the L7Cre mice. In the absence of Cre expression, the transgene is not produced. In the presence of Cre expression, the transgene will be “FLip-EXchanged” leading to expression of the transgene specifically in PCs.

### *In situ* hybridization

Fresh frozen 20µm thick-sections were prepared using a cryostat (Cryostar NX 70, Thermo Fisher Scientific, Ref.: 957000H) from brains of *Susd4* WT and KO mice at P0, P7 or P21. The probe sequence corresponded to the nucleotide residues 287-1064bp (encompassing exons 2-5) for mouse *Susd4* (NM_144796.4) cDNA. The riboprobes were used at a final concentration of 0.05µg/µL, and hybridization was done overnight at a temperature of 72°C. The anti-digoxigenin-AP antibody (for details see antibodies section above) was used at a dilution of 1:5000. Alkaline phosphatase detection was done using BCIP/NBT colorimetric revelation (Roche Life Science, Cat#11681451001).

### Behavioral Study

12-14 weeks old male mice were used in this study. They were housed in groups of 3-5 in standard conditions: 12h light/dark cycle, with *ad libitum* food and water access. Seven days before the beginning of behavioral test, mice were housed individually to limit inter-houses variability resulting from social relationships. All behavioral tests took place in the light cycle.

#### S.H.I.R.P.A. protocol

Mice performed a series of tests to ensure their general good health and motor performance and habituate them to being manipulated (Crawley, 2006). The test includes observation of appearance, spontaneous behavior, neurological reflexes, anxiety, motor coordination, balance rotarod and muscular strength tests and were performed within five days. Individuals presenting deficits during the S.H.I.R.P.A. protocol were not used for other behavioral tests.

#### Footprint analysis

The fore and hind paws of mice were dipped in magenta and cyan non-toxic paint, respectively. Mice were allowed to walk through a rectangular plastic tunnel (9cm W × 57cm L × 16cm H), whose floor was covered with a sheet of white paper. Habituation was done the day before the test. Five footsteps were considered for the analysis. Footprints were scanned and length measurements were made using ImageJ.

#### Rotarod

Mice were first habituated to the rotarod apparatus, three days before the acceleration test. The habituation protocol consists of 5min at 4 r.p.m. To evaluate the motor coordination, mice were placed on immobile rotarod cylinders, which ramped up from 0 to 45 rotations per minute in 10min. The timer was stopped when the mouse fell off the cylinder or did a whole turn with it. For a given session, this procedure was repeated three times per day separated by 60min. The session was repeated during five consecutive days.

## Whole-cell patch-clamp on acute cerebellar slices

Responses to parallel fiber (PF) and CF stimulation were recorded in PCs of the lobule VI in acute parasagittal and horizontal (long-term potentiation (LTP) experiments) cerebellar slices from *Susd4* KO juvenile (from P25 to P35) or adult (∼P60) mice. *Susd4* WT littermates were used as controls. Mice were anesthetized using isoflurane 4% and sacrificed by decapitation. The cerebellum was dissected in ice cold oxygenated (95% O_2_ and 5% CO_2_) Bicarbonate Buffered Solution (BBS) containing (in mM): NaCl 120, KCl 3, NaHCO_3_ 26, NaH_2_PO_4_ 1.25, CaCl_2_ 2, MgCl_2_ 1 and D(+)-glucose 35. 300µm-thick cerebellar slices were cut with a vibratome (Microm HM650V: Thermo Scientific Microm, Massachusetts, USA or 7000smz-2 Campden Instruments Ltd., UK) in slicing solution (in mM): N-Methyl-D-Glucamine 93, KCl 2.5, NaH_2_PO_4_ 1.2, NaHCO_3_ 30, HEPES 20, D(+)-Glucose 25, MgCl_2_ 10, sodium ascorbate 5, thiourea 2, sodium pyruvate 3, N-acetyl-cystein 1, kynurenic acid 1 and CaCl_2_ 0.5 (pH 7.3). Immediately after cutting, slices were allowed to briefly recover at 37°C in the oxygenated sucrose-based buffer (in mM): sucrose 230, KCl 2.5, NaHCO_3_ 26, NaH_2_PO_4_ 1.25, D(+)-glucose 25, CaCl_2_ 0.8 and MgCl_2_ 8. D-APV and minocycline at a final concentration of 50µM and 50nM, respectively, were added to the sucrose-based buffer. Slices were allowed to fully recover in bubbled BBS with 50Mm minocycline at 37°C for at least 40min before starting the experiment, then maintained at RT for a maximum time of 8h (from slicing time). Patch clamp borosilicate glass pipettes with 3-6MΩ resistance were filled with the following internal solutions:

1. Cesium metanesulfonate solution (CsMe solution, for EPSC elicited from CF and PF), containing (in mM) CsMeSO_3_ 135, NaCl 6, MgCl_2_ 1, HEPES 10, MgATP 4, Na_2_GTP 0.4, EGTA 1.5, QX314Cl 5, TEA 5 and biocytin 2.6 (pH 7.3).
2. CsMe S-solution (for delayed EPSC quanta events), containing (in mM): CsMeSO_3_ 140, MgCl_2_ 0.5, HEPES 10, MgATP 4, Na_2_GTP 0.5, BAPTA 10 and neurobiotin 1% (pH 7.35).
3. Potasium Gluconate solution (KGlu solution, for PF long-term plasticity), containing (in mM): K Gluconate 136, KCl 10, HEPES 10, MgCl_2_ 1, sucrose 16, MgATP 4 and Na_2_GTP 0.4 (pH 7.35).

Stimulation electrodes with ∼5 MΩ resistances were pulled from borosilicate glass pipettes and filled with BBS. Recordings were performed at room temperature on slices continuously perfused with oxygenated BBS. The experiment started at least 20min after the whole-cell configuration was established. The Digitimer DS3 (Digitimer Ltd) stimulator was used to elicit CF and PF and neuronal connectivity responses in PCs. Patch-clamp experiments were conducted in voltage clamp mode (except for the LTP and long-term depression (LTD) induction protocols that were made under current clamp mode) using a MultiClamp 700B amplifier (Molecular Devices, California, USA) and acquired using the freeware WinWCP written by John Dempster (
https://pureportal.strath.ac.uk/en/datasets/strathclyde-electrophysiology-software-winwcp-winedr). Series resistance was compensated by 60-100% and cells were discarded if significant changes were detected. Currents were low-pass filtered at 2.2kHz and digitized at 20kHz.

### CF and PF-EPSC experiments

To isolate the AMPARs current, the BBS was supplemented with (in mM) picrotoxin 0.1, D-AP5 10, CGP52432 0.001, JNJ16259685 0.002, DPCPX 0.0005 and AM251 0.001. CF and PF EPSCs were monitored at a holding potential of -10mV. During CF recordings, the stimulation electrode was placed in the granule cell layer below the clamped cell; CF-mediated responses were identified by the typical all-or-none response and strong depression displayed by the second response elicited during paired pulse stimulations (20Hz). The number of CFs innervating the recorded PC was estimated from the number of discrete CF-EPSC steps. PF stimulation was achieved by placing the stimulation electrode in the molecular layer at the minimum distance required to avoid direct stimulation of the dendritic tree of the recorded PC. The input-output curve was obtained by incrementally increasing the stimulation strength. Peak EPSC values for PF were obtained following averaging of three consecutive recordings, values for CF-EPSC correspond to the first recording. Short-term plasticity experiments were analyzed using a software written in Python by Antoine Valera (http://synaptiqs.wixsite.com/synaptiqs).

### PF-Long-term plasticity experiments

PCs were clamped at -60mV. Each PF-induced response was monitored by a test protocol of paired stimulation pulses (20Hz) applied every 20s. A baseline was established during 10min of paired-pulse stimulation in the voltage clamp configuration. After that, an induction protocol was applied in current-clamp mode with cells held at -60mV. During LTD induction, the PFs were stimulated with two pulses at high frequency (200Hz) and, after 100ms, the CF was stimulated with four pulses at high frequency (200Hz) repeated every 2 seconds for a period of 10min. During LTP induction, the PFs were stimulated with bursts of 15 pulses at high frequency (100 Hz) repeated every 3s for a period of 5min (Binda et al., 2016). Then, PCs were switched to the voltage clamp mode and paired stimulation pulses applied again, lasting 40min. All the data were normalized to the mean baseline. Long-term plasticity was analyzed with the software Igor Pro 6.05 (WaveMetrics INC, Oregon, USA).

### PF and CF delayed EPSC quanta events

were detected and analyzed using the software Clampfit 10.7 (Molecular Devices). PF- and CF-delayed EPSC quanta superposed events were discarded manually based on the waveform. A threshold of 10pA for minimal amplitude was used to select the CF events. 100 (PF) and 300 (CF) events for each neuron were studied by analyzing consecutive traces.

## High density microelectrode array (MEA) analysis of Purkinje cell spiking in acute cerebellar slices

Experiments were performed on acute cerebellar slices obtained from 3-6 months-old mice in artificial cerebrospinal fluid (ACSF) containing (in mM): NaCl 125, KCl 2.5, D(+)Glucose 25, NaHCO_3_ 25, NaH_2_PO_4_ 1.25, CaCl_2_ 2, and MgCl_2_ 1 and oxygenated (95% O_2_ and 5% CO_2_). Parasagittal slices (320µm) were cut at 30°C (Huang and Uusisaari, 2013) with a vibratome (7000smz-2, Campden Instruments Ltd.) at an advance speed of 0.03mm/s and vertical vibration set to 0.1 - 0.3µm. Slices were then transferred to a chamber filled with oxygenated ACSF at 37°C and allowed to recover for 1h before recordings.

For recording, the slices were placed over a high-density micro electrode array of 4096 electrodes (electrode size, 21 × 21µm; pitch, 42µm; 64 × 64 matrix; Biocam X, 3Brain, Wädenswil, Switzerland), and constantly perfused with oxygenated ACSF at 37C°. Extracellular activity was digitized at 17 kHz and data were analyzed with the Brainwave software (3Brain). The signal was filtered with a butterworth high-pass filter at 200 Hz, spikes were detected with a hard threshold set at -100µV, and unsupervised spike sorting was done by the software. We selected units with a firing rate between 15 and 100 spikes per second and we excluded units presenting more than 5% of refractory period violation (set to 3ms). Recordings were performed on two slices per animal, each slice containing between 20 and 200 active neurons, and results were then pooled for each animal.

To quantify the average variability in the firing rate, the coefficient of variation (CV) of the interspike interval (ISI) in seconds) was calculated as the ratio of the standard deviation (SD) of ISIs to the mean ISI of a given cell. To measure the firing pattern variability within a short period of two ISIs, CV2 was calculated [CV2 = 2|ISI_n+1_ − ISI_n_|/(ISI_n+1_ + ISI_n_)] (Holt and Douglas, 1996).

## Affinity-purification of SUSD4 interactors from synaptosome preparations

HEK293H (Gibco, Massachusetts, USA, Cat#11631-017) were maintained at 37°C in a humidified incubator with 5% CO_2_ in Dulbecco’s Modified Eagle’s Medium (DMEM; containing high glucose and glutamax, Life Technologies, Cat#31966047) supplemented with 10% fetal bovine serum (FBS, Gibco, Cat#16141-079), and 1% penicillin/streptomycin (Gibco, Cat#15140122). 10^6^ cells were plated per well in a 6-well plate and transfected 24 hours (h) after plating with the indicated plasmids (1µg plasmid DNA per well) using Lipofectamine 2000 (Invitrogen, Cat#11668-019) according to manufacturer’s instructions.

48h after transfection, cells were lysed and proteins were solubilized for 1h at 4°C under gentle rotation in lysis buffer (10mM Tris-HCl pH7.5, 10mM EDTA, 150mM NaCl, 1% Triton X100 (Tx; Sigma, Cat#x100), 0.1% SDS) supplemented with a protease inhibitor cocktail (1:100; Sigma, Cat#P8340) and MG132 (100µM; Sigma, Cat#C2211). Lysates were sonicated for 10 seconds, further solubilized for 1h at 4°C and clarified by centrifugation at 6000 r.p.m. during 8 minutes (min). Supernatants were collected, incubated with 5µg of rat monoclonal anti-HA antibody (for details see antibodies), together with 60µL of protein G-sepharose beads (Sigma; Cat#10003D) for 3h at 4°C, to coat the beads with the HA-tagged SUSD4 proteins. When SUSD4-GFP was expressed for affinity-pulldowns, GFP-Trap was done according to the instructions of GFP-Trap®_A (Chromotek, New York, USA, Cat#ABIN509397). Coated beads were washed 3 times with 1mL lysis buffer.

To prepare synaptosome fractions, cerebella from WT mice (P30) were homogenized at 4°C in 10 volumes (w/v) of 10mM Tris buffer (pH7.4) containing 0.32M sucrose and protease inhibitor cocktail (1:100). The resulting homogenate was centrifuged at 800g for 5min at 4°C to remove nuclei and cellular debris. Synaptosomal fractions were purified by centrifugation for 20min at 20000 r.p.m. (SW41Ti rotor) at 4°C using Percoll-sucrose density gradients (2-6-10-20%; v/v). Each fraction from the 10 - 20% interface was collected and washed in 10mL of a 5mM HEPES buffer pH 7.4 (NaOH) containing 140mM NaCl, 3mM KCl, 1.2mM MgSO_4_, 1.2mM CaCl_2_, 1mM NaH_2_PO_4_, 5mM NaHCO_3_ and 10mM D(+)-Glucose by centrifugation. The suspension was immediately centrifuged at 10000g at 4°C for 10min. Synaptosomes in the pellet were resuspended in 100µL of lysis buffer (10mM Tris-HCl pH7.5, 10mM EDTA, 150mM NaCl, 1% Tx) supplemented with a protease inhibitor cocktail (1:100) and MG132 (100µM). Lysates were sonicated for 10 seconds, and further incubated for 1h at 4°C. HA-SUSD4, GFP-SUSD4 or its control GFP coated beads were then incubated with the synaptosomal lysates for 3h at 4°C. Beads were washed three times with lysis buffer supplemented with 0.1% SDS. Bound proteins were eluted for 10min at 75°C using Laemmli buffer (160mM Tris ph6.8, 4% SDS, 20% glycérol, 0.008% BBP) with 5% β-mercaptoethanol before SDS-PAGE followed by western blotting or mass spectrometry.

## Co-Immunoprecipitation experiments in HEK293 cells

10^6^ HEK293H cells were plated per well in 6-well plates and transfected 24h after plating with the indicated plasmids (1.6µg plasmid SEPGLUA2 per well, using a molar ratio of 2:1 SEPGLUA2:other plasmid) using Lipofectamine 2000 according to manufacturer’s instructions. For anti-HA pull downs, proteins from HEK293 cell lysates were solubilized in lysis buffer (1M Tris-HCl pH8, 10mM EDTA, 1,5M NaCl, 1% Tergitol ^TM.^ (sigma; Cat#NP40), 2% Na azide, 10% SDS and 10% Na deoxycholate) supplemented with a protease inhibitor cocktail (1:100) and MG132 (1%). Then, lysates were sonicated for 15s, further clarified by a centrifugation at 14000 r.p.m. for 10min. Supernatants were collected and incubated with Dynabeads protein G (life technologies, Cat#10004D) and 28.8µg of rat monoclonal anti-HA antibody (for details see antibodies) under gentle rotation for 1h at 4°C. Precipitates were washed three times in lysis buffer and then eluted by boiling (65°C) the beads 15min in sample buffer (made from sample buffer 2X concentrate, Sigma, Cat#S3401) before SDS-PAGE. For SEPGLUA2 pull downs, 48h after transfection, cells were washed twice in 1X PBS, lysed with 200µL of lysis buffer (50mM Tris-HCl pH8 and 1% Tx) supplemented with a protease inhibitor cocktail (1:100) and MG132 (50µM), scraped, sonicated 3 × 5 seconds, and proteins were further solubilized for 30min at 4°C under rotation. Lysates were clarified by centrifugation at 14000 r.p.m. for 10min at 4°C. Supernatants (inputs) were collected and incubated with G-protein Dynabeads (ThermoFisher Scientific, Cat#10004D), previously linked to mouse anti-GFP antibody (for details see antibodies section), under gentle rotation for 1h at 4°C, to coat the beads with the SEP-tagged GLUA2 proteins and interactors. Using a magnet, coated beads were washed five times in lysis buffer and bound proteins were then eluted by boiling for 15min at 65°C in 1X sample buffer before SDS-PAGE and western blot analysis for detection of HA-SUSD4 and GLUA2.

## Mass spectrometry analysis

Proteins were separated by SDS-PAGE on 10% polyacrylamide gels (Mini-PROTEAN® TGX™ Precast Gels, Bio-Rad, Hercules USA) and stained with Protein Staining Solution (Euromedex, Souffelweyersheim France). Gel lanes were cut into five pieces and destained with 50mM triethylammonium bicarbonate (TEABC) and three washes in 100% acetonitrile. Proteins were digested in-gel using trypsin (1.2µg/band, Gold, Promega, Madison USA), as previously described (Thouvenot et al., 2008). Digest products were dehydrated in a vacuum centrifuge.

### Nano-flow liquid chromatography coupled to tandem mass spectrometry (NanoLC-MS/MS)

Peptides, resuspended in 3µL formic acid (0.1%, buffer A), were loaded onto a 15cm reversed phase column (75mm inner diameter, Acclaim Pepmap 100® C18, Thermo Fisher Scientific) and separated with an Ultimate 3000 RSLC system (Thermo Fisher Scientific) coupled to a Q Exactive Plus (Thermo Fisher Scientific) *via* a nano-electrospray source, using a 120min gradient of 5 to 40% of buffer B (80% ACN, 0.1% formic acid) and a flow rate of 300nL/min.

MS/MS analyses were performed in a data-dependent mode. Full scans (375 - 1,500m/z) were acquired in the Orbitrap mass analyzer with a 70000 resolution at 200m/z. For the full scans, 3 × 10^6^ ions were accumulated within a maximum injection time of 60ms and detected in the Orbitrap analyzer. The twelve most intense ions with charge states ≥ 2 were sequentially isolated to a target value of 1 × 10^5^ with a maximum injection time of 45ms and fragmented by HCD (Higher-energy collisional dissociation) in the collision cell (normalized collision energy of 28%) and detected in the Orbitrap analyzer at 17500 resolution.

### MS/MS data analysis

Raw spectra were processed using the MaxQuant environment ((Cox and Mann, 2008), v.1.5.5.1) and Andromeda for database search (Cox et al., 2011). The MS/MS spectra were matched against the UniProt Reference proteome (Proteome ID UP000000589) of *Mus musculus* (release 2017_03; http://www.uniprot.org) and 250 frequently observed contaminants (MaxQuant contaminants database) as well as reversed sequences of all entries. The following settings were applied for database interrogation: mass tolerance of 7ppm (MS) and 0.5 Th (MS/MS), trypsin/P enzyme specificity, up to two missed cleavages allowed, only peptides with at least seven amino acids in length considered, and Oxidation (Met) and acetylation (protein N-term) as variable modifications. The “match between runs” (MBR) feature was allowed, with a matching time window of 0.7min. FDR was set at 0.01 for peptides and proteins.

A representative protein ID in each protein group was automatically selected using an in-house bioinformatics tool (leading v2.1). First, proteins with the most numerous identified peptides are isolated in a “match group” (proteins from the “Protein IDs” column with the maximum number of “peptides counts”). For the match groups where more than one protein ID is present after filtering, the best annotated protein in UniProtKB (reviewed entries rather than automatic ones), highest evidence for protein existence, most annotated protein according to the number of Gene Ontology Annotations (GOA Mouse version 151) is defined as the “leading” protein. Only proteins identified with a minimum of two unique peptides, without MS/MS in control immunoprecipitation and exhibiting more than 4-fold enrichment (assessed by spectral count ratio) in Sushi domain-containing protein 4 (SUSD4) immunoprecipitation, *vs* control immunoprecipitation, in the two biological replicates, were considered as potential partners of SUSD4 (**Table 1**).

### Gene Ontology analysis

The statistically enriched gene ontology (GO) categories for the 28 candidate proteins were determined by Cytoscape (v3.6) plugin ClueGO v2.5.3 (Bindea et al., 2009). The molecular function category was considered (release 18.12.2018, https://www.ebi.ac.uk/GOA), except evidences inferred from electronic annotations. Terms are selected by different filter criteria from the ontology source: 3-8 GO level intervals, minimum of 4 genes per GO term and 10% of associated genes/term. A two-sided hypergeometric test for enrichment analysis (Benjamini-Hochberg standard correction used for multiple testing) was applied against the whole identified protein as reference set. Other predefined settings were used. Each node representing a specific GO term is color-coded based on enrichment significance (p-value). Edge thickness represents the calculated score (kappa, κ) to determine the association strength between the terms.

## Chemical LTD and GLUA2 surface biotinylation assay in cerebellar acute slices

300 µm-thick parasagittal cerebellar slices were obtained from P31-P69 WT and *Susd4* KO mice following the same protocol described before (Patch-clamp section). Slices were incubated for 2h at 37°C in oxygenated BBS with or without proteasome (50µM MG132 in DMSO,) and lysosomal (100µg/mL leupeptine in water, Sigma, Cat#11034626001) inhibitors. Chemical LTD was induced by incubating the slices for 5min at 37°C in BBS containing 50mM K^+^ and 10µM glutamate (diluted in HCl), followed by a recovery period in BBS for 30min at 37°C all under oxygenation; in presence or not of inhibitors. Control slices were incubated in parallel in BBS solution containing HCl. Slices were then homogenized in lysis buffer, containing: 50mM Tris-HCl, 150mM NaCl, 0.1% SDS, 0.02% Na Azide, 0.5% Na Deoxycholate, 1% NP-40 and protease inhibitor cocktail (1:100). Homogenates were incubated 45min at 4°C, then sonicated and centrifuged at 14000 r.p.m. for 10min at 4°C. Supernatants were then heated at 65°C in 2X sample buffer (Sigma, Cat#S3401) prior to western blot analysis for detection of GLUA2 and GLURδ2. For GLUA2 surface biotinylation assay, cerebellar slices (obtained from mice aged between P27-P61) were treated as above. After a recovery period of 30min at 37°C in BBS, slices were incubated in a biotinylation solution (ThermoFisher Scientific, EZ-Link™ Sulfo-NHS-SS-Biotin, Cat#A39258, 0,125mg/mL) for 30min on ice without oxygen. Slices were finally washed three times for 10min in PBS pH7.4 at 4°C and then homogenized in lysis buffer, containing: 50mM Tris-HCl pH8, 150mM NaCl, 0.1% SDS, 0.02% Na Azide, 0.5% Na Deoxycholate, 1% NP-40 and protease inhibitor cocktail (1:100). Homogenates were incubated 45min at 4°C, then sonicated and centrifuged at 14000 r.p.m. for 10min at 4°C. Supernatants (inputs) were collected and incubated with Dynabeads MyOne Streptavidin C1 (Thermo Fisher Scientific, Cat#65001) under gentle rotation overnight at 4°C. Using a magnet, beads were washed five times in lysis buffer and biotinylated proteins were then eluted by boiling for 15min at 65°C in 1X sample buffer before SDS-PAGE and western blot analysis for detection of GLUA2.

## Immunocytochemistry

### Labeling of primary hippocampal neurons

Hippocampi were dissected from E18 mice embryos and dissociated. 1.2×10^5^ neurons were plated onto 18 mm diameter glass cover-slips precoated with 80µg/mL poly-L-ornithine (Sigma, Cat#P3655) and maintained at 37°C in a 5% CO_2_ humidified incubator in neurobasal medium (Gibco, Cat#21103049) supplemented with 2% B-27 supplement (Gibco, Cat#17504044) and 2mM Glutamax (Gibco, Cat#35050-038). Fresh culture medium (neurobasal medium supplemented with 2% B-27, 2mM L-glutamine (Gibco, Cat#A2916801) and 5% horse serum (Gibco, Cat#26050088) was added every week for maintenance of the neuronal cultures.

Hippocampal neurons at days *in vitro* 13 (DIV13) were transfected using Lipofectamine 2000 and 0.5µg plasmid DNA per well. After transfection, neurons were maintained in the incubator for 24h, then fixed with 100% methanol for 10min at -20°C. After rinsing with PBS, non-specific binding sites were blocked using PBS containing 4% donkey serum (DS, Abcam, Cat#ab7475) and 0.2% Tx Primary and secondary antibodies were diluted in PBS 1% DS / 0.2% Tx and incubated 1h at room temperature. Three washes in PBS 0.2% Tx were performed before and after each antibody incubation. Nuclear counterstaining was performed with Hoechst 33342 (Sigma, Cat#14533) for 15min at room temperature.

### Labeling of primary cerebellar mixed cultures

Cerebellar mixed cultures were prepared from P0 tg/0 “B6.129-Tg(Pcp2-cre)2Mpin/J” (Stock Number: 004146, outbred, C57Bl/6J background) mouse cerebella and were dissected and dissociated according to previously published protocol (Tabata et al., 2000). Neurons were seeded at a density of 5×106 cells/mL. Mixed cerebellar cultures were transduced at DIV3 using a Cre-dependent AAV construct that express HA-tagged SUSD4 and soluble GFP (2µL of AAV2-hSYN-DIO-HA-SUSD4-2A-eGFP-WPRE at 4,1.10^12 GC/mL or control AAV2-hSYN-DIO-eGFP-WPRE at 5.10^12 GC/mL). At DIV17, neurons were fixed with 4% PFA in PBS1X for 30min at room temperature. After rinsing with PBS, non-specific binding sites were blocked using PBS containing 4% DS and 0.2% Tx. Primary and secondary antibodies were diluted in PBS 1% DS and 0.2% Tx and incubated one hour at room temperature. Three PBS 0.2% Tx washes were performed before and after each antibody incubation. Nuclear counterstaining was performed with Hoechst 33342 for 15min at room temperature.

## Immunohistochemistry

### Labeling of brain sections

30µm-thick parasagittal brain sections were obtained using a freezing microtome (SM2010R, Leica) and brains obtained after intracardiac perfusion with 4% PFA in PBS solution of mice sedated with 100mg/kg pentobarbital sodium. Sections were then washed three times for 5min in PBS, then blocked with PBS 4% DS for 30min. The primary antibodies were diluted in PBS, 1% DS, 1% Tx. The sections were incubated in the primary antibody solution overnight at 4°C and then washed three times for 5min in PBS 1% Tx. Sections were incubated in the secondary antibody, diluted in PBS 1% DS 1% Tx solution, for 1h at room temperature. The sections were then incubated for 15min at room temperature with the nuclear marker Hoechst 33342 in PBS 0.2% Tx. Finally, the sections were washed three times for 5min in PBS 1% Tx, recovered in PBS and mounted with Prolong Gold (Thermo Fisher Scientific, Cat#P36934) between microscope slides and coverslips (Menzel-gläser, Brunswick, Germany, Cat#15165252).

## RT-PCR and quantitative RT-PCR

For standard RT-PCR, total RNA was isolated from the cortex, cerebellum and brainstem of 2-month-old *Susd4* KO mice and WT control littermates, using the RNeasy mini kit (Qiagen, Venlo, Netherlands, Cat#74104). Equivalent amounts of total RNA (100 ng) were reverse-transcribed according to the protocol of SuperScript® VILO™ cDNA Synthesis kit (Life Technologies, California, USA, Cat#11754-250) as stated by manufacturer’s instructions. The primers used were forward 5’ TGT TAC TGC TCG TCA TCC TGG 3’ and reverse 5’ GAG AGT CCC CTC TGC ACT TGG 3’. PCR was performed with an annealing temperature of 61°C, for 39 cycles, using the manufacturer’s instructions (*Taq* polymerase; New England Biolabs, Massachusetts, USA, Cat#M0273S). Quantitative PCR was performed using the TaqMan universal master mix II with UNG (applied biosystems, Cat# 4440038) and the following TaqMan probes: *Rpl13a* (#4331182_Mm01612986_gH) and *Susd4* (#4331182_Mm01312134_m1).

## Western Blot analysis

After samples were mixed with sample buffer, proteins were resolved by electrophoresis on a 4-12% NuPAGE Bis-Tris-Gel according to Invitrogen protocols, then electrotransferred using TransBlot DS Semi-dry transfer Cell or TransBlot Turbo transfer system (Bio-Rad) to PVDF membrane (Immobilon-P transfer membrane, Millipore, Cat#IPVH00010). Membranes were blocked in PBS supplemented with Tween 0.2% (PBST) and non-fat milk 5% and incubated with primary antibodies in PBST-milk 5%. After washing three times in PBST, membranes were incubated with Horseradish Peroxidase-conjugated secondary antibodies in PBST-milk 5%. Membranes were finally washed three times and bound antibodies were revealed using Immobilon Western (Millipore, Cat#WBKLS) or Western Femto Maximum Sensitivity (Thermo Fisher Scientific, Cat#34095) or SuperSignal West Dura (Thermo Fisher Scientific, Cat#34075) or ECL Western Blotting substrate (Thermo Fisher Scientific, Cat#32209) chemiluminescent solutions and images acquired on a Fusion FX7 system (Vilber Lourmat, Île-de-France, France). Quantitation of Western blots was performed using the ImageJ software on raw images under non-saturating conditions. Band intensities of proteins of interest were obtained after manually selecting a rectangular region around the band. The signal intensity of the band of interest was then normalized to the signal intensity of the corresponding βACTIN (used as a loading control). For quantifications of immunoprecipitation experiments, input intensities were normalized to βACTIN, and then the intensities of immunoprecipitated protein bands were normalized to the normalized inputs.

## Image acquisition and quantification

*In situ* hybridization images were acquired using an Axio Zoom. V16 (Zeiss, Oberkochen, Germany) microscope equipped with a digital camera (AxioCam HRm) using a 10x objective (pixel size 0.650µm).

Immunofluorescence image stacks were acquired using a confocal microscope (SP5, Leica), using a 63x objective (1,4NA, oil immersion, pixel size: 57nm for cell culture imaging, pixel size: 228nm for 63x; 76nm, 57nm, 45nm for higher magnifications for *in vivo* imaging). The pinhole aperture was set to 1 Airy Unit and a z-step of 200 nm was used. Laser intensity and photomultiplier tube (PMT) gain was set so as to occupy the full dynamic range of the detector. Images were acquired in 16-bit range. Immunofluorescence images and image stacks from figure 1C, 1D and 4F were acquired using a Zeiss LSM 980 Confocal with an Airyscan detector (v2.0), using a 63x objective (1,4NA, oil immersion, pixel size: 43nm, z-step of 150nm).

Deconvolution was performed for the VGLUT1 images with Huygens 4.1 software (Scientific Volume Imaging) using Maximum Likelihood Estimation algorithm from Matlab. 40 iterations were applied in classical mode, background intensity was averaged from the voxels with lowest intensity, and signal to noise ratio values were set to a value of 25.

VGLUT1 and VGLUT2 puncta were analyzed using the Matlab software and a homemade code source (Dr. Andréa Dumoulin). The number, area and intensity of puncta were quantified using the mask of each puncta generated by the Multidimensional Image analysis software (MIA) from Metamorph® (Molecular Devices). For each animal, puncta parameters were measured from four equidistant images within a 35-image stack at 160 nm interval, acquired from three different lobules (n=12).

The software ImageJ was used to measure the total area of a cerebellar section from images of staining obtained with the nuclear marker Hoechst. The extension of the molecular layer was measured using images of the anti-CABP staining. Nine parasagittal sections were analyzed per animal. The data presented correspond to the mean per animal.

## Statistical analysis

Data from all experiments were imported in Prism (GraphPad Software, California, USA) for statistical analysis, except for electrophysiology data that were imported to Igor Pro 6.05 (WaveMetrics INC) for statistical analysis.

In the case of two column analyses of means, the differences between the two groups were assessed using two-tailed Student’s t-test. Normality of populations were assessed using D’Agostino & Pearson, Shapiro-Wilk and Kolmogorov-Smirnov normality tests. When groups did not fit the normal distribution, the non-parametric Mann-Whitney test was used. For the rotarod behavioral test (two variables, genotype and trial), two-way repeated measures ANOVA followed by Bonferroni post hoc test was performed. The two-tailed Student’s one sample t-test (when normality criterion was met) or the two-tailed Wilcoxon Signed Rank Test was used to compare ratios to a null hypothesis of 1 for biochemical experiments or 100 for long-term plasticity (Fay, 2013). Differences in cumulative probability were assessed with the Kolmogorov-Smirnov distribution test, and differences in distribution were tested using the Chi-squared test.

